# Specificity and promiscuity of JAK recruitment regulates pleiotropy of cytokine-receptor signaling

**DOI:** 10.1101/2023.10.04.560821

**Authors:** Eyal Zoler, Thomas Meyer, Junel Sotolongo Bellón, Boyue Sun, Jacob Piehler, Gideon Schreiber

## Abstract

Promiscuous binding of different Janus kinases (JAKs) to class I/II cytokine receptors has been reported, yet its role in signaling is unclear. To systematically explore JAK pairing in type I interferon (IFN-I) signaling, we generated an artificial IFN-I receptor (AIR) by replacing the extracellular domains of IFNAR1 and IFNAR2 with anti mEGFP and mCherry nanobodies. The heterodimeric AIR restored near-native IFN-I activity, while the homomeric variant of IFNAR2 (AIR-dR2) initiated much weaker signaling despite harboring docking sites for signal transducer and activator of transcription (STAT) proteins. AIR-dR1 was signaling inactive, yet, pulldown uncovered its ICD to bind both TYK2 and JAK1. To further investigate the roles of JAKs on the receptors, knockout (KO) JAK1, JAK2, TYK2, and JAK2/TYK2 were generated. JAK1 KO led to complete loss of IFN-I signaling, which was partially restored by TYK2 overexpression. TYK2 KO cells retained partial activity, which was elevated by JAK1 overexpression, suggesting both JAKs to partially substitute each other. Conversely, JAK2 KO only moderately impacted the biological activity of IFN-Is, even in JAK2/TYK2 KO cells. Live cell micropatterning confirmed promiscuous binding of JAK1, JAK2 and TYK2 to IFNAR1 and IFNAR2, in line with an AlphaFold model that shows JAKs interchangeability on IFNAR ICDs. Similar promiscuity of JAK binding was observed for TPOR and GHR but not EPOR, accompanied by different downstream signaling activity. The competitive binding of JAKs to cytokine receptors together with the highly diverse absolute and relative JAK expression levels can account for cell type-dependent signaling pleiotropy observed for cytokine receptors.

**One Sentence Summary:** Promiscuous and interchangeable binding of JAKs to cytokine receptors enables cell type-specific pleiotropic signaling.

## INTRODUCTION

Type I interferons (IFN-I) are helical cytokines with vital functions in initiating innate and adaptive immune responses against a wide range of pathogens (*1*). Additionally, they act as crucial regulators of tumor immunity and contribute to developing autoimmune diseases. These IFN-Is are secreted proteins that promote antiviral (AV) activity in virtually every nucleated cell in vertebrates, while antiproliferative (AP), and immunomodulatory activities are cell type specific (*2–4*). The human family of IFN-I consists of 17 members. These include 13 subtypes of IFN-α with high sequence and structural similarity (*5–7*), as well as IFN-β, IFN-κ, IFN-ω, and IFN-ε (*8*, *9*). All these members share a common cell surface receptor, comprised of the subunits IFN-α receptor 1 (IFNAR1) and IFN-α receptor 2 (IFNAR2) (*3*, *10*).

IFNAR1 and IFNAR2 are single spanning transmembrane proteins classified as class 2 helical cytokine receptors. IFNAR2 is the high-affinity receptor, while IFNAR1 binds IFN-I with lower affinity (*11*). The intracellular domains (ICDs) of human IFNAR1 and IFNAR2 are 100 and 251 amino acids long, respectively. IFNAR1 and IFNAR2 activate signaling via Janus family tyrosine kinases (JAKs), which non-covalently associate with membrane-proximal segments of the ICDs characterized by “Box1” and “Box2” motifs (*12*). The “Box1” motif is characterized by its proline-rich nature, while the “Box2” motif typically includes a negatively-charged residue followed by several hydrophobic residues. Although these two motifs show low conservation among different receptors, they are essential for ensuring proper JAK activity across various JAK-receptor pairs (*12*, *13*). IFNAR1 and IFNAR2 have been shown to recruit TYK2 and JAK1, respectively, via their Box1/Box2 motifs. While the Box1 motif of IFNARs have minimal impact on JAK1 or TYK2 binding, they play a significant role in their activation (*14*). The signaling cascade of IFN-I is initiated by binding to IFNAR1 and IFNAR2 (*11*). This binding brings the two receptor subunits close together, facilitating their associated Janus kinases (JAKs) cross-phosphorylation, which subsequently activates the JAK-STAT signaling pathway within the cell (*15*, *16*). Once activated, the JAKs phosphorylate various other proteins, including signal transducers and activators of transcription (STAT) proteins, particularly STAT1 and STAT2 (*17–19*) that are recruited via docking sites mostly located on the ICD of IFNAR2 (*14*). Phosphorylated STAT1 and STAT2 are core components of transcription factors, which are responsible for upregulating expression of interferon-stimulated genes (ISGs) (*20–22*). ISGs play a role in blocking viral entry, translation, and replication. The IFN-I signaling cascade is a complex and tightly regulated process, playing a crucial role in the body’s defense against viral infections and preventing viral entry and replication. In turn, IFN-Is and components of their signaling cascade are preferential targets of inhibition by the viruses (*23–26*). As a master regulator of antiviral defense, IFN-I signaling emerged as a paradigm of signaling plasticity that has been shown to depend on IFN-receptor interaction dynamics, cell surface receptor concentration and negative feedback regulators (*1*, *3*, *27–31*). Next to robust antiviral activity elicited in largely all cell types, more complex responses such cell death or immunomodulatory activities are strongly dependent of the cellular context (*30*). However, the role of JAK expression and potential alternate usage of JAKs by IFNAR in regulating IFN-I signaling plasticity has so far not been systematically explored. The JAK family comprises four members: JAK1, JAK2, JAK3, and TYK2, all constitutively associated with cytokine receptors ICDs. JAK proteins consist of four distinct structural parts: an N-terminal FERM (Band 4.1, Ezrin, Radixin, Moesin-homology) domain, an SH2 domain, a pseudokinase (PK) domain, and a C-terminal catalytic (tyrosine kinase) domain (TK) (*32*). Extensive research has established that individual JAK family members physically interact with various cytokine receptors. The FERM and SH2 domains of JAK proteins have been identified as vital components in mediating the JAK-receptor interaction. Through co-purification studies and crystallization of the FERM-SH2 region of JAKs in complex with their respective associated receptors (TYK2-IFNAR1, JAK1-IFNLR1, and JAK2-LEP-R/JAK2-EPOR), it has been observed that the FERM and SH2 domains of JAKs associate to form a module that is responsible for binding to the receptors (*33–35*). The FERM and SH2 domains are essential for engaging with the cytoplasmic tails of the receptors containing the Box1 and Box2 motives, respectively. Moreover, several studies have demonstrated a critical role of JAK binding to stabilize receptor cell surface expression, most prominently for IFNAR1, which critically depends on the association of TYK2 (*36*, *37*).

To study the structure/function relations of the ICDs, a synthetic cytokine receptor system was recently developed, in which the receptor extracellular domains were replaced with nanobodies binding mEGFP and mCherry fused to the transmembrane and intracellular domains of the desired receptor. Homo- and heterodimerization of such synthetic receptors can be flexibly controlled by corresponding homo- or heteromeric mEGFP/mCherry variants that bind to the nanobodies (*38*). The capabilities of this framework has been validated for diverse receptor systems (*39*, *40*) including murine IFNAR (*41*) . Here, we deployed this approach to design an artificial IFN-I receptor (AIR) to study the role of JAKs in human IFN-I signaling. We found that in the absence of TYK2 and JAK2, significant IFN-I signaling is possible via JAK1/JAK1 homodimer due to promiscuous JAK binding of IFNAR1. Our study provides insights into the roles of JAK1, JAK2, and TYK2 in IFN-I signaling and their changeability on the receptors. The findings support the hypothesis of multiple kinase binding and highlight the significance of JAK1 homodimers. Additionally, we uncover the impact of JAK1, JAK2 and TYK2 on receptor trafficking and reveal broader implications for the regulation of cytokine receptor signaling in general.

## RESULTS

### An Artificial IFN-I Receptor (AIR) Signals Similar to Native IFNAR

AIR was generated by utilizing nanobodies against GFP (α-GFP) and mCherry (α-mCherry) to replace the ectodomains (ECD) of hIFNAR1 (AIR-R1) and hIFNAR2 (AIR-R2), respectively (Fig. 1A). The AIR ligand (AIR-L) heterodimerizing IFNAR1 and IFNAR2 consists of mEGFP and mCherry (GFP-mCherry) connected by a five-amino-acid linker. The ligand was expressed in *E. coli* and purified by immobilized metal ion chromatography, sumo protease cleavage, anion exchange chromatography, and size exclusion chromatography (Fig. S1A-B). For quantifying AIR cell surface expression, we used flow cytometry (FACS), using the ALFA-tag fused to the N-terminus of the receptors. We found that the surface expression of AIR was ∼10-fold higher than the wild-type receptors (Fig. 1B). As overexpression of the native receptor elevates surface levels by only 2-3 fold (*42*), this suggest that the ECDs contribute to the tight control of IFNAR cell surface expression. Upon treating cells expressing AIR-R1 and AIR-R2 with AIR-L, near native activation of the JAK-STAT pathway was observed. Western Blot (WB) revealed robust phosphorylation of STAT1, 2, 3, 5, and 6, comparable to the wild-type receptors treated with saturation concentration of IFN-β (Fig. 1C). The AIR system exhibited robust ISG induction after 8 and 16 hours as determined by qPCR measuring both robust and tunable gene(*6*) (Fig. 1D), similar to that observed by IFN-β treatment. To further correlate transcriptional activity, we purified RNA from HeLa cells after 16 hours of treatment with IFN-β and the AIR ligand to determine activation of gene expression. Fig. 1E shows a comparison of upregulated genes between IFN-β and AIR as determined by MARSseq, showing a similar overall pattern of expression. This was further verified by pathway analysis using the GSEA software (*43*) (Fig. S1C). The IFN-I and IFN-II pathways were the major upregulated ones, exhibiting a high adjusted p-value (padj). Antiviral (AV) potency against vesicular stomatitis virus (VSV) and encephalomyocarditis virus (EMCV) showed EC50 values nearly identical between AIR and IFN-β (Fig. 1F and G).

**Figure 1:**
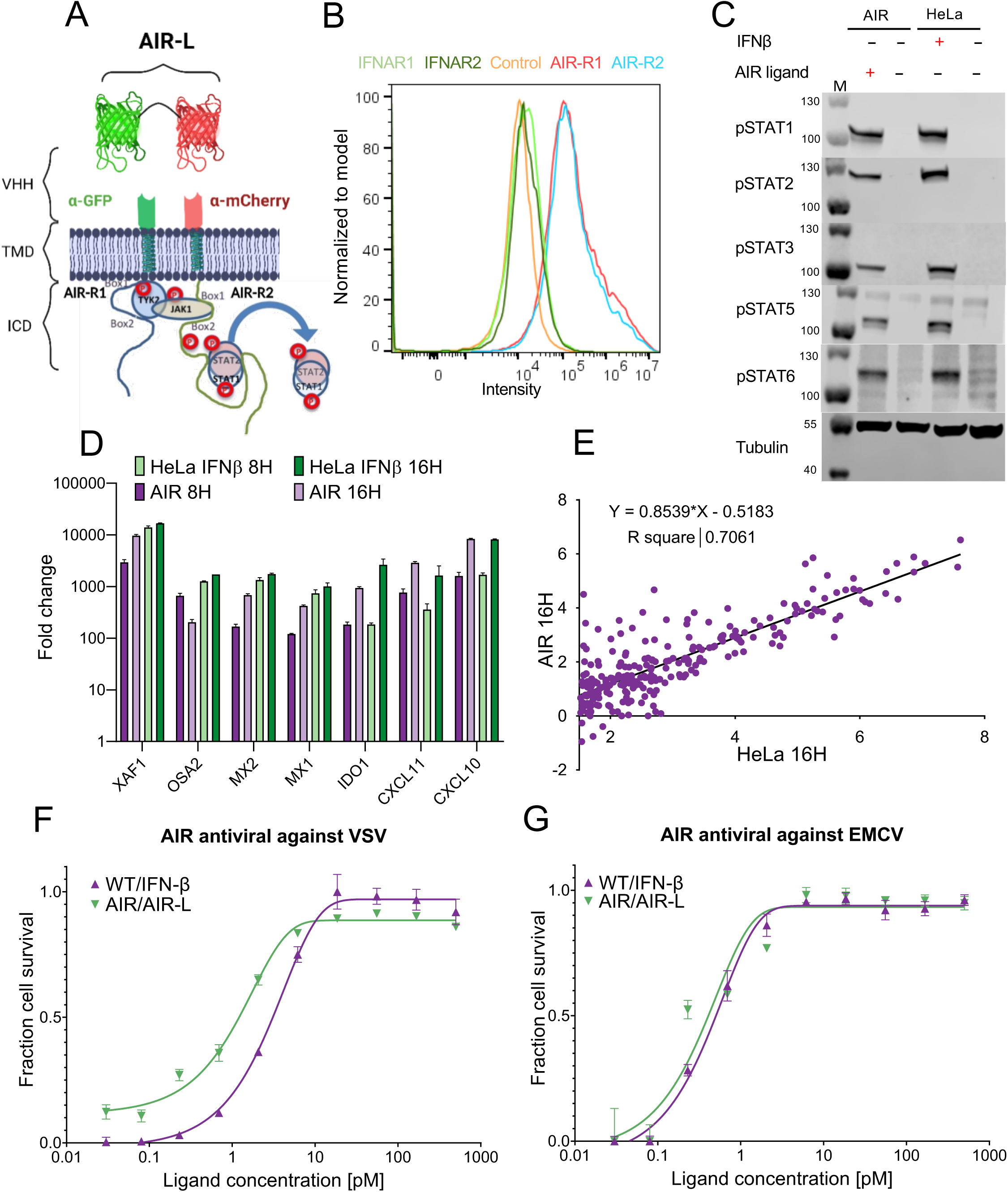
Characterization of the human artificial interferon receptor (AIR) and its activation of the JAK-STAT Pathway. (**A**) A schematic illustration depicting the concept of the AIR system, which consists of anti-GFP (α-GFP) and anti-mCherry (α-mCherry) nanobodies fused to the transmembrane and intracellular domains of hIFNAR1 and hIFNAR2, respectively. (**B**) Flow cytometry 48 hrs after transient transfection of ALFA-IFNAR1, ALFA-IFNAR2, ALFA-AIR-R1, and ALFA-AIR-R2, respectively. Non-permeabilized cells were stained using a labeled anti-ALFA-tag nanobody (n = 3 independent experiments). (**C**) WT HeLa cells and HeLa transfected with AIR-R1 and AIR-R2 (AIR) were treated with IFN-β or AIR-L for 30 min and analyzed by Western blot for phosphorylated STAT proteins. Tubulin is a loading control. M, mass markers. Data are representative of n = 2 independent experiments. (**D**) Gene expression in response to treatment with IFN-β or AIR-L for 6 and 16 hrs. The data represent the fold change compared to untreated cells and normalized to HPRT1. XAF1, MX1, MX2, and OAS2 are IFN-I induced robust genes; IDO1, CXCL10 and CXCL11 are IFN-I tunable genes. The results show the average and SD of n = 3 independent experiments per group. (**E**) Cells were treated for 16 hrs with 2 nM IFN-β or AIR ligand and subjected to MARSseq. The data presented are of genes whose expression level increased by 2-fold or higher by IFN-β in WT HeLa cells. The x-axis is log(2) of HeLa treated with IFN-β, and the y-axis is log(2) of AIR treated with the AIR ligand. (**F-G**) Antiviral Activity against VSV and EMCV for HeLa cells and HeLa cells 48 hours post transient transfection with AIR. Cells were treated with IFN-β or AIR-L for 4 hrs before infection with the VSV or EMCV for 18 hrs or 20 hrs, respectively. Cells were stained with crystal violet for cell viability.

### Homodimeric AIR Complexes maintain residual activity

To test the relevance of receptor homodimerization, we developed two other versions of the AIR receptors: an IFNAR1 homodimer (AIR-dR1) and an IFNAR2 homodimer (AIR-dR2) (Fig. 2A). For AIR-dR1 the ligand dimerizes two IFNAR1 ICDs, while for AIR-dR2 the ligand dimerizes two IFNAR2 ICDs. While AIR-dR1 did not promote STAT phosphorylation, AIR-dR2 phosphorylated both STAT1 and STAT2, albeit to low levels (Fig. 2B). In line with the observed STAT phosphorylation, AIR-dR2 induced expression of interferon-stimulated gene, which was not observed for AIR-dR1 (Fig. 2C). Finally, we evaluated AV potency of dR1 and dR2 against VSV and EMCV. AIR-dR1 did not show any antiviral activity, whereas AIR-dR2 exhibited a low level of activity, which correlated with lower levels of phosphorylated STATs and reduced gene induction in comparison to HeLa cells treated with IFN-β (Fig. 2D, E). The lack of signaling activity observed for the AIR-dR1 homodimer is expected as only the ICD of IFNAR2 were shown to bind and activate STAT proteins (*14*). To further verify that signaling inactivity of AIR-dR1 is caused by lack of JAK binding, we performed pull-down of AIR-dR1 combined with mass-spectrometry (Table S1). In addition to TYK2, we found surprisingly high levels of JAK1 bound to dR1, but as expected no STAT pulldown. With this in mind, AIR-dR2 homodimer could signal either through JAK1/TYK2 binding or JAK1/JAK1 binding.

**Figure 2.**
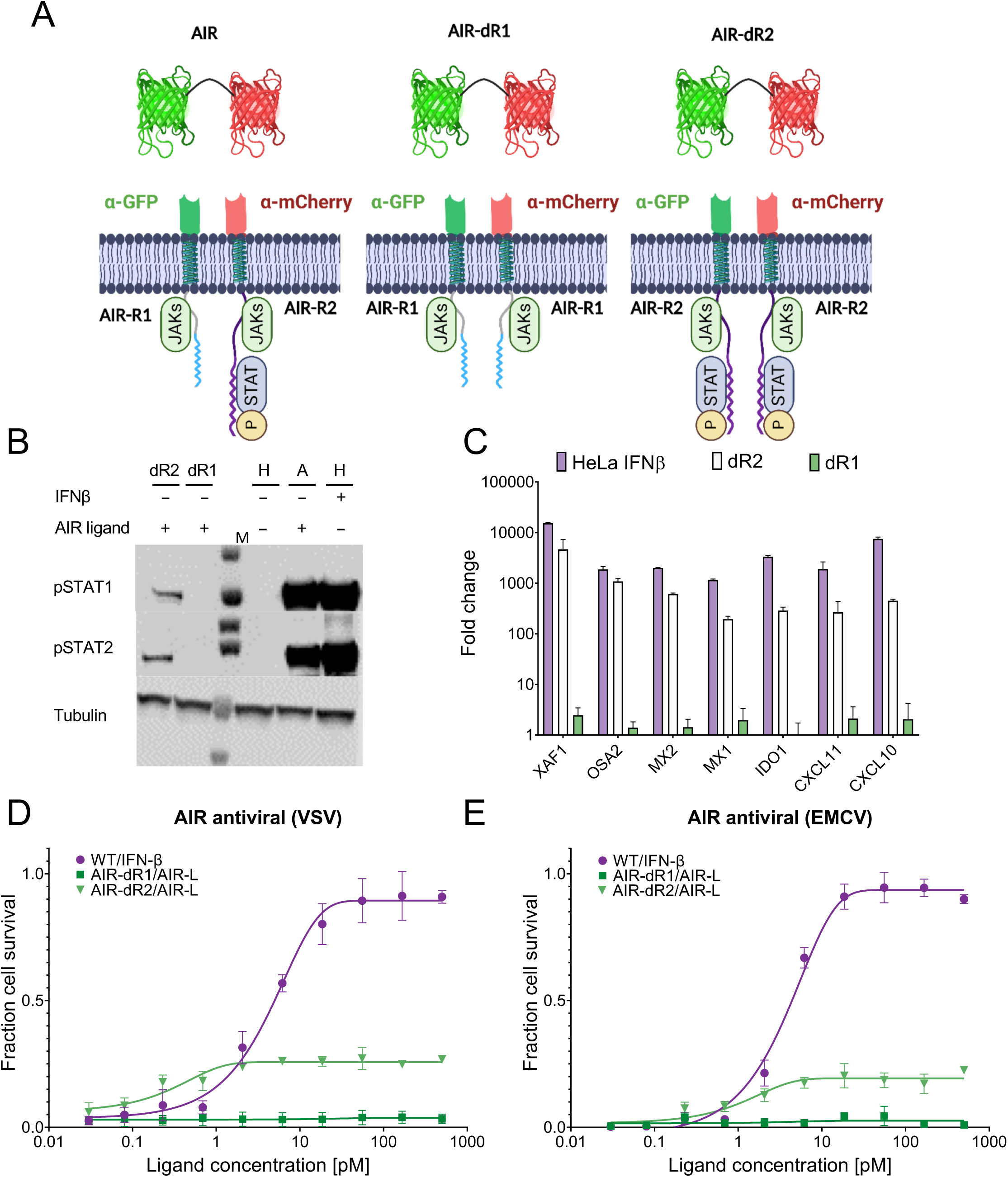
Activation of IFN-I signaling by homo-dimer AIR. (**A**) illustrates the composition of two homodimer versions of the AIR system: AIR-dR1 (homodimer IFNAR1) and AIR-dR2 (homodimer IFNAR2), in addition to the heterodimer AIR (AIR-FL). (**B**) HeLa cells (H) and HeLa Transfected with AIR-FL (A), AIR-dR1 (dR1) and AIR-dR2 (dR2) constructs. Cells were treated with IFN-β or AIR-L for 30 min and analyzed by Western blot for phosphorylated STAT proteins. Tubulin is a loading control. M, mass markers. Data are representative of n = 2 independent experiments. (**C**) Gene expression in response to treatment with IFN-β or AIR-L for 6 and 16 hrs. The data represent the fold change compared to untreated cells and normalized to HPRT1. XAF1, MX1, MX2, and OAS2 are IFN-I robust genes; IDO1, CXCL10 and CXCL11 are IFN-I tunable genes. The results show the average and SD of n = 3 independent experiments per group. (**D, E**) Antiviral activity against VSV (**D**) and EMCV (E) for WT HeLa cells and HeLa cells 48 hours post transient transfection of AIR. Cells were treated with IFN-β or AIR-L for 4 hrs before infection with the VSV or EMCV for 18 hrs or 20 hrs, respectively. Cells were stained with crystal violet for cell viability.

### Both JAK1 and TYK2 are not absolutely Essential for Type I Interferon Signaling

To further investigate the role of JAK family members in IFN-I signaling, we first performed individual knockout experiments for each JAK protein. As reported in a previous study (*14*), the KO of JAK1 leads to complete inhibition of STAT phosphorylation upon treatment with IFN-β, along with elimination of IFN-I gene induction and biological activity (Fig. 3 and Fig. S2). Reconstituting JAK1 rescued the IFN-I activity of the KO cells, including the AV activity (Fig. 3C, D), and phosphorylation of STAT1 and STAT2 (Fig. 3A). What was not expected is that overexpression of TYK2 in JAK1 KO background resulted in partial, although low, activation of pSTAT1 and pSTAT2 (Fig. 3A). This suggests that overexpressed TYK2 can bind the ICD of IFNAR2, and is activated by TYK2/TYK2 cross-phosphorylation.

**Figure 3.**
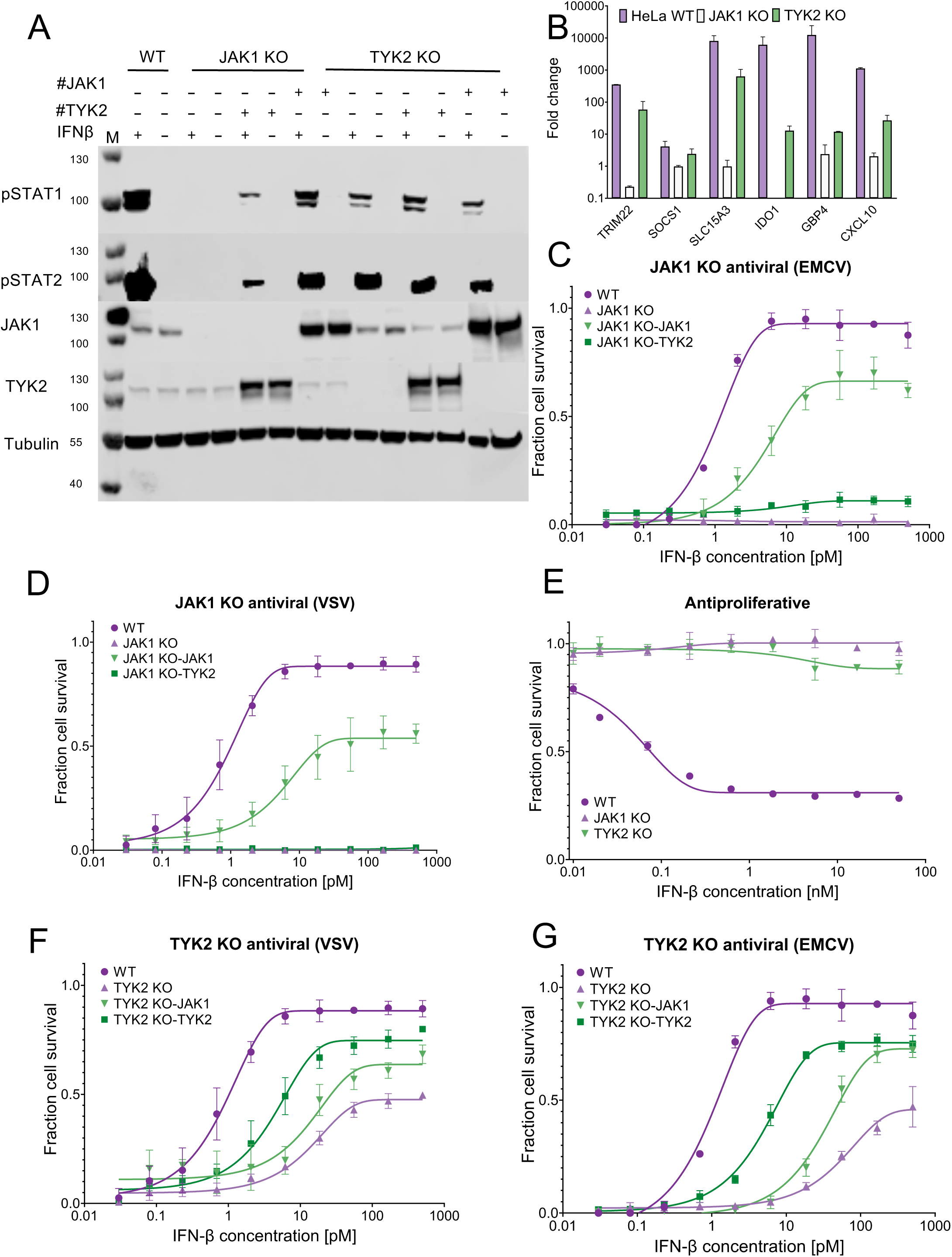
IFN-I signaling in JAK1 and TYK2 KO cells. **(A)** STAT phosphorylation of HeLa cells (WT), HeLa JAK1 KO and HeLa TYK2 KO cells after 30 min of treatment with 2 nM IFN-β, relative to non-treated cells. #JAK1 and #TYK2 designate cells transiently transfected with JAK1 or TYK2 for 48 hours prior to treatment. Tubulin is a loading control. M, mass markers. Data are representative of n = 2 independent experiments. (**B**) Gene expression in response to treatment with IFN-β or AIR ligand for 16 hrs. The data represent the fold change compared to untreated cells and normalized to HPRT1. TRIM22, SLC15A3, and, OAS2 are IFN-I robust genes; IDO1, CXCL10 and GBP4 are IFN-I tunable genes. The results show the average and SD of n = 3 independent experiments per group. (**C**) Antiproliferative activity, HeLa, JAK1 KO, and TYK2 KO cells upon treatment with IFN-β for 96 h before staining with crystal violet for cell viability. Data are representative of n = 3 to 5 independent experiments. Error bars represent 95% CI. **(D-G**) Antiviral activity against VSV and EMCV for HeLa, JAK1 KO, and TYK2 KO cells treated with IFN-β for 4 hrs before infection with the VSV or EMCV for 18 hrs or 20 hrs, respectively. Cells were stained with crystal violet for cell viability. Data are representative of n = 3 independent experiments. Error bars represent 95% CI.

The contribution of TYK2 KO towards IFN-I activity has been less conclusive. In both mice and humans, some reports suggested it to be essential (*41*, *44*) while others showed residual IFN-I induced activity, particularly STAT1 activation (*45–47*). Here, we generated HeLa TYK2 KO cells using CRISPR/Cas9 (Fig. 3A). In line with the later, treatment with IFN-β showed STAT1 and STAT2 activation (albeit weaker than in WT cells – Fig. 3A and S2). It should be noted that pSTAT4 and 5 levels are much more reduced that pSTAT1 and 2 levels upon TYK2 KO. Following gene expression upon IFN-β treatment showed significant increase in expression of robust genes (TRIM22, SOCS1, SLC15A3), while expression of tunable genes (IDO1, GBP4, CXCL10) were only marginally induced (Fig. 3B). In line with these results, TYK2 KO abolished the AP activity upon IFNβ treatment (Fig. 3C), while partial AV activity was maintained both against VSV and EMCV (Fig. 3F, G). These findings collectively affirm the vital and indispensable role of TYK2 for uncompromised activation of the IFN-I pathway. However, significant residual activity in the absence of TYK2 was maintained, suggesting that another JAK can functionally replace it. Intriguingly, we observed that overexpression of JAK1 in TYK2 KO cells further potentiated the activation of the pathway (Fig. 3A, F-G), underscoring its significance and suggesting that JAK1 can replace TYK2 in IFNAR signaling.

### JAK2 is Relevant for full IFN-I Antiviral Activity

The ambiguous roles of JAKs in the activation of IFNAR prompted us to investigate the involvement of JAK2, which has so far not been implicated in IFN-I signaling (JAK3 is not expressed on HeLa cells). The potency IFN-I to initiate an AV state against VSV in the background of JAK2 KO was similar to WT cells. However, when infecting the cells with EMCV, the level of protection did not reach 100% at high concentrations of IFN-β (Fig. 4A, B). This suggests a role for JAK2 in IFN-I signaling. The EC50 of antiproliferative activity of IFN-β was not affected by JAK2 KO, but the fraction of surviving cells increased from 0.3 (WT) to 0.5 (Fig. 4C). These results suggest a lower potency of IFN-I in JAK2 KO cells. To complement the KO data, we overexpressed JAK2 in HeLa cells and observed that this resulted in complete protection against both VSV and EMCV (Fig. 4E, F). To evaluate whether this protection involves the IFNAR receptors, we overexpressed JAK2 in the background of IFNAR1/2 KO cells. Here, we observed a 30% decrease in protection, suggesting that part of the antiviral protection conferred by JAK2 overexpression is associated with the type I IFN pathway.

**Figure 4.**
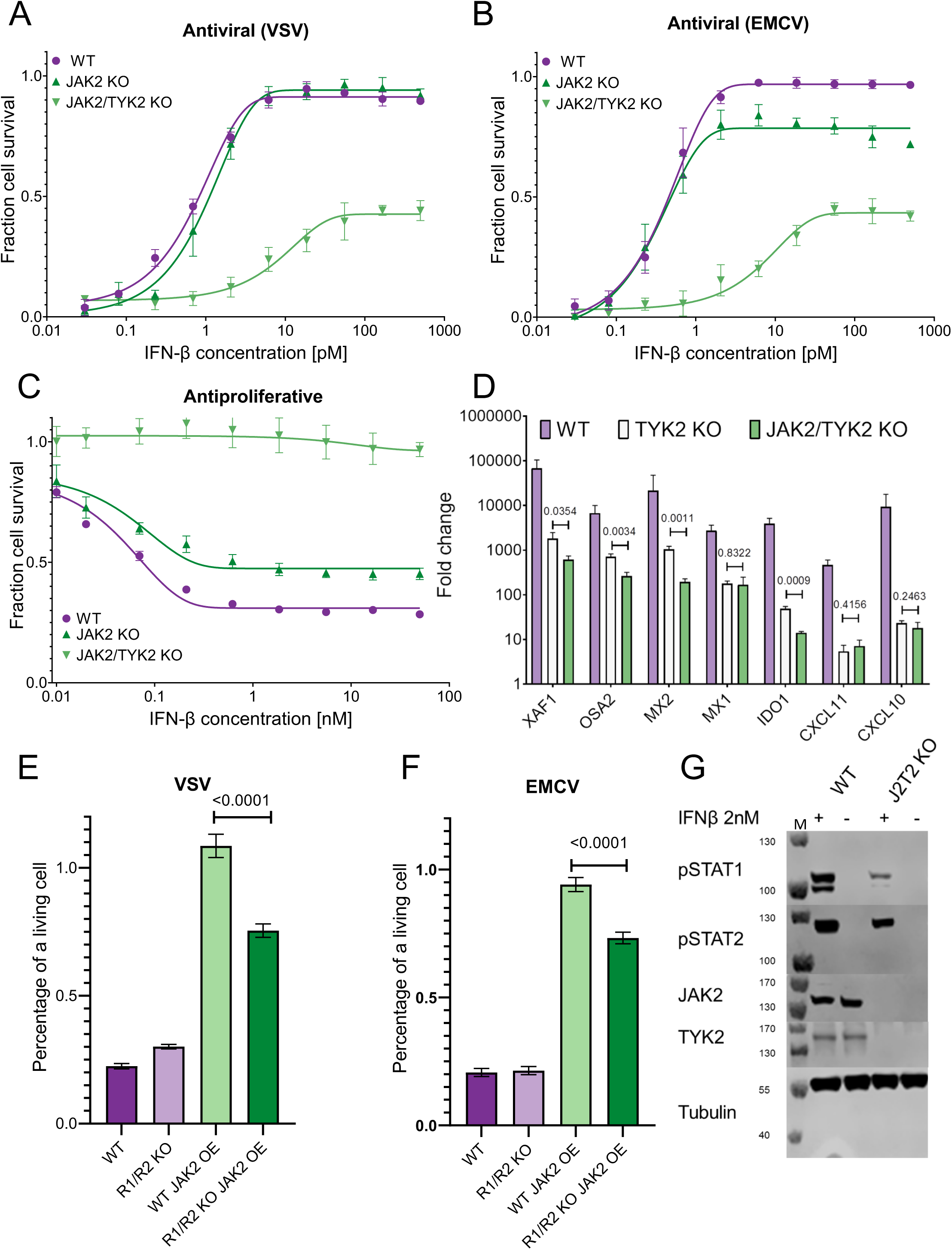
Effect of JAK2 Knockout on IFN-I signaling. (**A-B**) Antiviral activity against VSV and EMCV for HeLa, JAK2 KO, and JAK2/TYK2 KO cells treated with IFN-β for 4 hrs before infection with the VSV or EMCV for 18 hrs or 20 hrs, respectively. Cells were stained with crystal violet for viability. (**C**) Antiproliferative activity of HeLa, JAK2 KO, and JAK2/TYK2 KO cells following treatment with IFN-β for 96 hrs before staining with crystal violet for cell viability. Data are representative of n = 3 to 5 independent experiments. Error bars represent 95% CI. (**D**) Gene expression in response to treatment with IFN-β for 18 hrs. The data represent the fold change compared to untreated cells and normalized to HPRT1. XAF1, MX1, MX2, and OAS2 are IFN-I robust genes; IDO1, CXCL10 and CXCL11 are IFN-I tunable genes. The results show the average and SD of n = 3 independent experiments per group **(E-F**) Fraction of surviving cells after infection with VSV and EMCV. HeLa (WT), IFNAR1/2 KO (R1/R2 KO), and HeLa and IFNAR1/2 KO cells were transiently transfected for 48 hours with JAK2 (WT JAK2 OE and R1/R2 KO JAK2 OE) and then infected with VSV or EMCV for 18 hrs or 20 hrs, respectively. Cells were stained with crystal violet for cell viability. Data are representative of n = 3 to 5±SD. (**G**) HeLa (WT) and JAK2/TYK2 KO cells were treated with 2 nM IFN-β for 30 min and analyzed by Western blotting for phosphorylated STAT proteins, JAK2 and TYK2. Tubulin is a loading control. M, mass markers. Data are representative of n = 2 independent experiments.

To verify that JAK1 homodimer formation is a possible route for IFN-I signaling, we created a double knockout of JAK2 and TYK2 (J2T2-dKO). This approach allowed us to isolate the effects of other JAK proteins in the pathway. As with TYK2 KO cells, IFN-β did induce partial STAT1 and STAT2 activation in the J2T2-dKO cells (Fig. 4G), as well as partial AV protection (Fig. 4A, B). In line with the TYK2 KO, the dKO did not induce AP activity (Fig. 4C). However, gene induction was somewhat lower in the double KO cells in comparison to TYK2 KO alone (Fig 4D). Overall, our findings suggest that the IFNAR1/IFNAR2 heterodimer can signal via JAK1/JAK1 homodimers, though with much lower efficiency as compared to JAK1/TYK2. Several members of the GP130 family are assumed to signal via JAK1/JAK1 homodimers and the activity of JAK1 homodimers have been furthermore supported by recent structures of the full-length JAK1 in an artificial dimer of the IFNLR1 ICD (*16*, *48*) Our observations suggest biological relevance of the JAK1/JAK1 homodimers in a heterodimeric receptor that potentially modulate its biological activity.

### Cell Surface Receptor Levels and Signaling Complexes Depend on the JAK identity

To further explore the role of different JAKs in IFN-I signaling we quantified the formation of IFNAR signaling complexes at the plasma membrane. IFNAR1 and IFNAR2 cell surface levels are tightly regulated, with few hundreds to few thousand copies being accessible to the ligand, depending of the cell type(*49*, *50*). JAKs play an important role in regulating receptor trafficking, with TYK2 binding being implicated in preventing endocytosis of IFNAR1 and thus stabilizing its cell surface expression (*36*) (*51*). We therefore explored receptor cell surface expression in JAK KO cells by FACS. As expected, TYK2 KO resulted in lower IFNAR1 surface expression levels, while JAK1 KO had no effect (Fig. 5A). Complementary, IFNAR2 surface expression levels were significantly decreased in JAK1 KO cells, but not in TYK2 KO cells (Fig. 5B and S3). Surprisingly, IFNAR1 expression levels were similarly reduced by JAK2 KO as by TYK2 KO, while IFNAR2 expression was not affected. No further changes in IFNAR cell surface expression were observed in the J2/T2-dKOs cells. These observations highlight that IFNAR1 trafficking is more sensitive to JAK availability than IFNAR2, and that beyond TYK2, at least also JAK2 is involved in stabilizing IFNAR1 cell surface expression.

**Figure 5.**
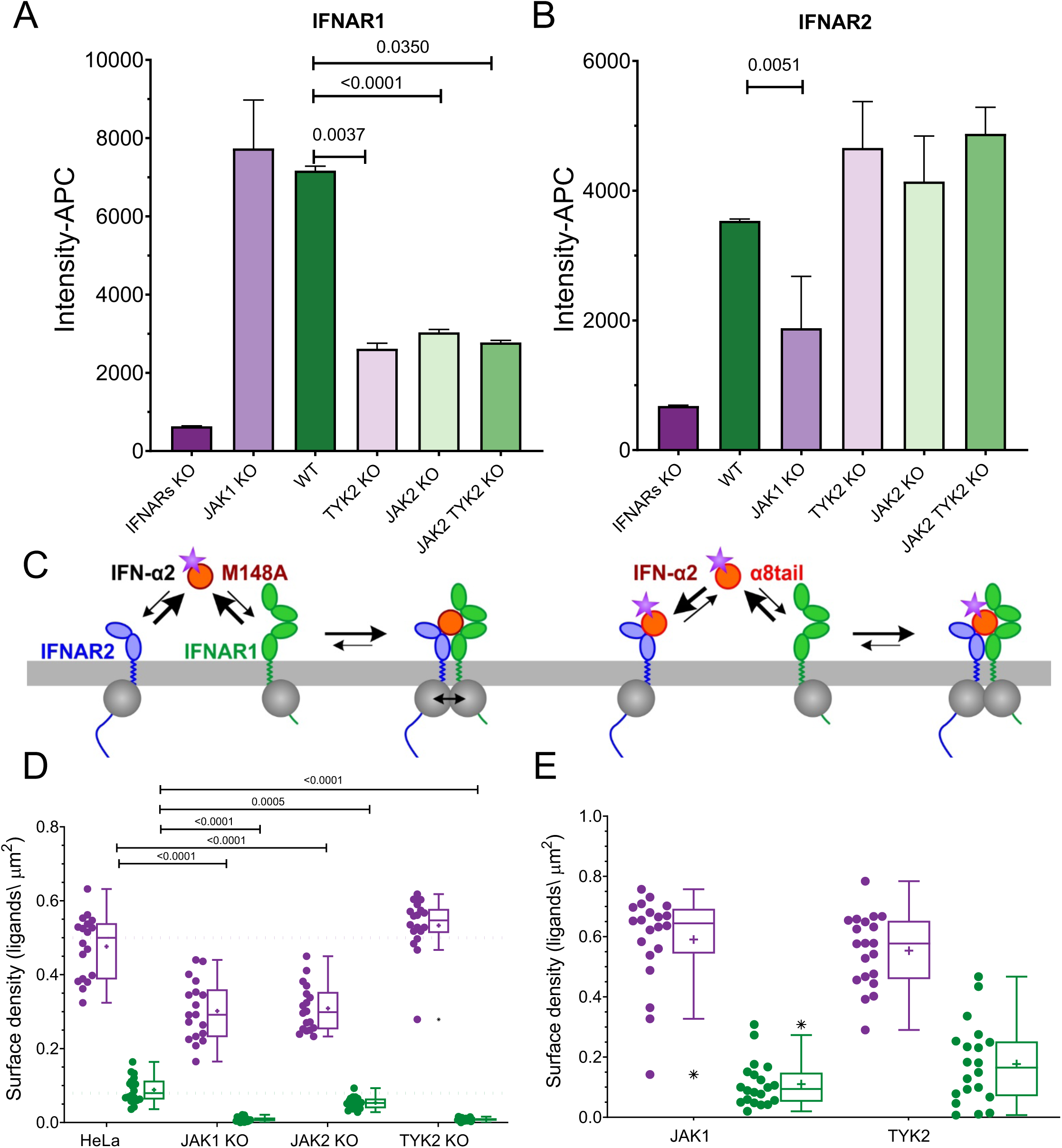
Cell surface receptor expression is regulated by JAKs. **(A-B**) Quantification of receptor cell surface expression levels in HeLa WT and KO cells by flow cytometry using antibodies against IFNAR1 (**A**) and IFNAR2 (**B**). Mean±SD from n = 3 to 5 independent experiments. **(C-E**) Formation of signaling complexes at the plasma membrane probed by quantifying IFN-α2 binding to endogenous receptor. (**C**) Cartoon of the assay: Binding of IFN-α2 with reduced IFNAR2 binding affinity (M148A, left) can only be detected upon efficient recruitment of IFNAR1, thus reporting receptor expression and productive interactions between the associated JAKs. The IFN-α2 variant α8tail (right) binds with very high affinity to IFNAR2 and therefore binds independently of IFNAR1. (**D**) Comparison of IFN-α2 M148A (green) and α8tail (purple) binding to WT and JAK KO HeLa cells. (**E**) IFN-α2 M148A and α8tail binding to TYK2 KO HeLa cell overexpressing JAK1 and TYK2, respectively.

To quantify how JAK KO affects the assembly of signaling complexes in the plasma membrane, we turned to single molecule localization microscopy (SMLM). To this end, we performed ligand binding assays using site-specifically fluorescent-labeled IFN-α2. We used the IFN-α2 mutant M148A, which binds IFNAR2 with ∼50-fold reduced affinity and thus requires simultaneous interaction with IFNAR1 for efficient binding (Fig. 5C). However, ternary complex assembly and thus IFN-α2 M148A binding also depends on the interactions between the JAKs PK domains, which has been shown to be important for JAK activation (*16*, *48*, *52*, *53*). Binding levels of IFN-α2 M148A therefore report on the overall capability to form signaling complexes in the plasma membrane (*52*). As a control, we used the IFN-α2 variant α8tail, which binds IFNAR2 with ∼20-fold increased affinity as compared to WT IFN-α2, and thus reports the cell surface density of IFNAR2. Representative binding experiments with IFN-α2 M148A and IFN-α2 α8tail binding to WT HeLa cells imaged by total internal reflection fluorescence (TIRF) microscopy are shown in Supplementary Movie S1. IFN-α2 binding to the surface was quantified as the density of localized fluorescent molecules. As expected, binding of IFN-α2 M148A significantly decreased in TYK2 KO (Fig. 5D), in line with the strongly reduced IFNAR1 cell surface expression level. By contrast, IFN-α2 α8tail binding was not affected, in line with the unchanged level of IFNAR2. In JAK1 KO cells, IFN-α2 α8tail densities were also reduced, as expected from the reduced IFNAR2 expression levels (Fig. 5D). However, IFN-α2 M148A binding was even more severely reduced, which can be explained by the lack of JAK1 also affecting the intracellular receptor interactions. In JAK2 KO cells, we found much less reduced binding of IFN-α2 M148A as compared to TYK2 KO cells, despite similar loss in IFNAR1 cell surface expression. This is in line with the interpretation that the presence of TYK2 still ensures optimum interaction with JAK1 in the signaling complex, which enhances receptor dimerization and downstream signaling. Overexpression of JAK1 and TYK2, respectively, in TYK2 knockout cells (Fig. 5E) yielded similarly increased binding of IFN-α2 M148A, suggesting that IFNAR1 cell surface expression and recruitment into the ternary complex can be rescued by JAK1 (Fig 5E). These results corroborate that IFNAR1 and IFNAR2 can engage different JAKs, yet with different efficiency.

### IFNAR1 and IFNAR2 Bind JAKs with Different Affinities

We therefore speculated that IFNAR1 and IFNAR2 bind different JAK members, yet with different affinities. To quantify binding affinities in the native context, we turned to live cell micropatterning (*54*). For this purpose, IFNAR subunits fused to an extracellular HaloTag and mTagBFP (HaloTag-mTagBFP-IFNAR) were expressed as bait proteins in HeLa cells and cultured on micropatterned substrates presenting the HaloTag ligand (HTL, Fig. 6A). This allowed for the capture and spatial reorganization of HaloTag-mTagBFP-IFNARs in the live cell plasma membrane. JAK interactions were probed by co-expressing mEGFP-tagged FERM/SH2 domains of different JAKs (JAK-FS), which are responsible for receptor binding (*36*). JAK-FS interacting with IFNARs enrich the micropatterns based on their binding affinity (Fig. 6B). The relative binding affinity was estimated by comparing the contrast of the micropatterns observed in the BFP and GFP channels using TIRF microscopy (Fig. 6C-F). In the case of IFNAR2, the contrast was significantly stronger for JAK1 compared to JAK2 and TYK2, suggesting a preferential binding of JAK1. However, in the JAK1 KO cells, the contrast for JAK2 increased by 173%, and for TYK2 by 70% (Fig. 6F), indicating that the other JAKs bind IFNAR2 as well, albeit at a lower affinity relative to JAK1 (Fig. 6D-F). Upon patterning IFNAR1, the highest contrast was observed for TYK2, while JAK1 and JAK2 gave lower, similar contrasts. Interestingly, the level of binding of JAK1 did not increase in the TYK2 KO cells, suggesting it is already saturated in the WT cells, which may be a result of the overexpression of the JAK1-FS domain (Fig. 6C-E). These results highlight a substantial promiscuity of IFNAR1 and IFNAR2 to bind different JAKs. In line with this observation, AlphaFold (*55*) predicted very similar modes of binding of JAK1, JAK2 and TYK2 FS domains in complex with the corresponding Box1/Box2 fragments of IFNAR1 and IFNAR2 (Fig. 7A).

**Figure 6.**
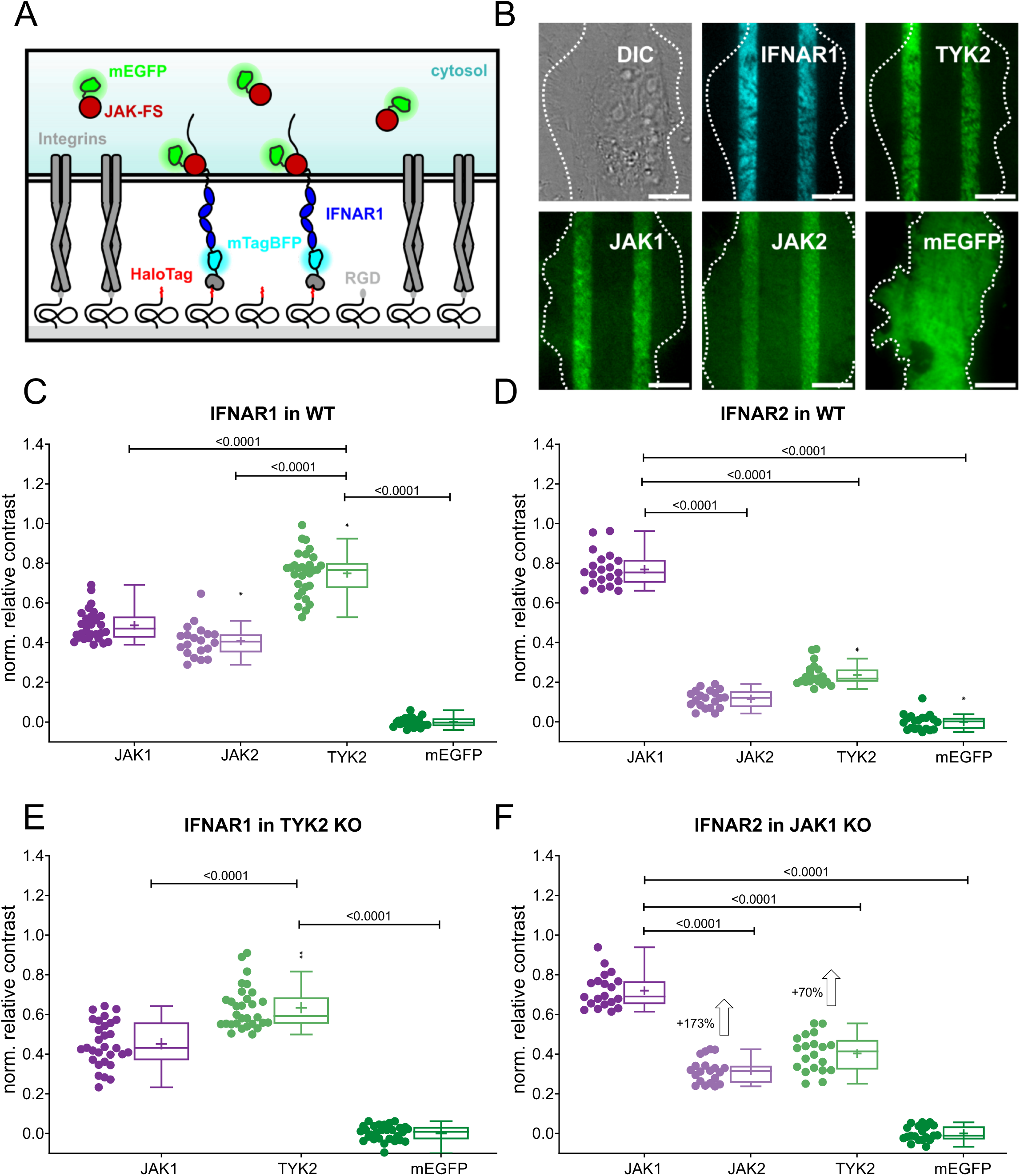
JAK1, JAK2 and TYK2 binding to IFNAR1 and IFNAR2 in live cells. **A**) Cartoon of live cell micropatterning to probe JAK binding to IFNAR1 in the plasma membrane. **B**) Representative experiments with micropatterned IFNAR1 (cyan) interacting with the mEGFP-tagged FERM/SH2 domain of different JAKs and mEGFP as negative control (green). Scale bar: 10 µm. C-F) Comparison of the contrast in receptor micropatterns observed for different JAK FERM/SH2 domains. C) IFNAR1-JAK interaction in WT HeLa cells. D) IFNAR2-JAK interaction in WT HeLa cells. E) IFNAR1-JAK interaction in HeLa TYK2 KO cells. F) IFNAR2-JAK interaction in HeLa JAK1 KO cells. Boxplots indicate the data distribution of the second and third quartile (box), median (line), mean (square), 1.5× IQR (whiskers), and minimum/maximum (x). Each data point in C), D), E), and F) represents the analysis from one cell. Statistical significances were determined by Kolmogorov-Smirnov analysis.

**Figure 7.**
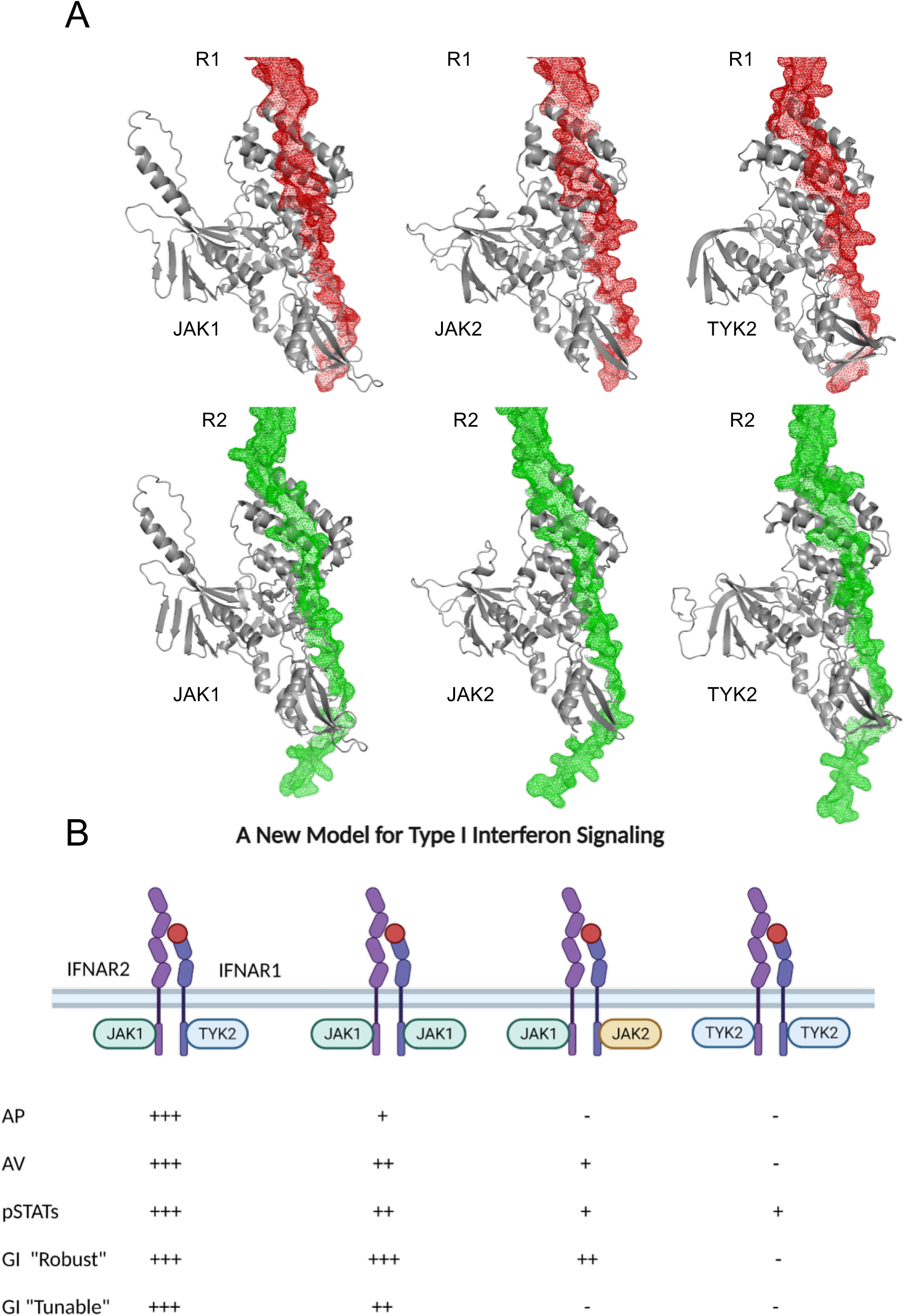
Modeling JAK binding to the ICDs of IFNAR1 and 2 using AlphaFold. **(B)** AlphaFold models of the FERM and SH2 domain of the given JAKs (JAK1 34-544, JAK2 37-482 and, TYK2 26-529) and the intracellular domain of IFNAR1 (437–507) and IFNAR2 (244–315). (**B**) Summary of activity of IFN-β signaling with different JAKs binding the ICDs (+++ 100% activation, ++ ∼50% activation, + weak activation, - not active).

### Promiscuity of JAKs Binding is a Common Feature in Cytokine Receptors

Taken together, our results clearly implicate that IFNAR1 and IFNAR2 each can recruit different JAK members with surprising promiscuity, yet optimum signaling activity requires JAK1 and TYK2. Our data suggest that rather than receptor binding, receptor trafficking and/or efficiency in cross-talk between different JAKs in the signaling complex are responsible for such differential activity. Promiscuity of JAK binding has been reported for several cytokine receptors (*56–59*), but its role in differentially regulating downstream signaling has hardly been appreciated (*59–61*). We propose that differential JAK recruitment may be a common principle, which may be overlooked due to a lack of suitable in situ receptor binding assays. To test this hypothesis, we turned to the homodimeric thrombopoietin receptor (TPOR), which has been reported to signal via JAK2 and to a lesser extent via TYK2. To test TPOR promiscuity toward JAKs, we first probed binding of JAK1, JAK2, TYK2 and JAK3 by live cell micropatterning with TPOR fused to HaloTag and mTagBFP (HaloTag-mTagBFP-TPOR) in WT HeLa cells. Interestingly, this receptor shows almost equally strong binding for JAK1, JAK2, and TYK2, as seen from the relative contrast generated by binding, while binding of JAK3 was significantly weaker (Fig. 8A, B). We further analyzed the stability of JAK-TPOR complexes using fluorescence recovery after photobleaching (FRAP). Since micropatterned TPOR is immobilized in the micropatterns, the recovery curve reflects the kinetics of photobleached JAK-FS being exchanged by cytosolic, unbleached JAK-FS (Fig. 8C), which is typically dominated by the dissociation of the TPOR/JAK complex. By fitting the recovery curve with a mono-exponential function, the lifetime of JAK-TPOR complexes were determined. This analysis resulted in very similar complex stabilities for all four members of the JAKs family (Fig. 8C), corroborating that TPOR can efficiently recruit JAK1, JAK2, JAK3, and TYK2.

**Figure 8.**
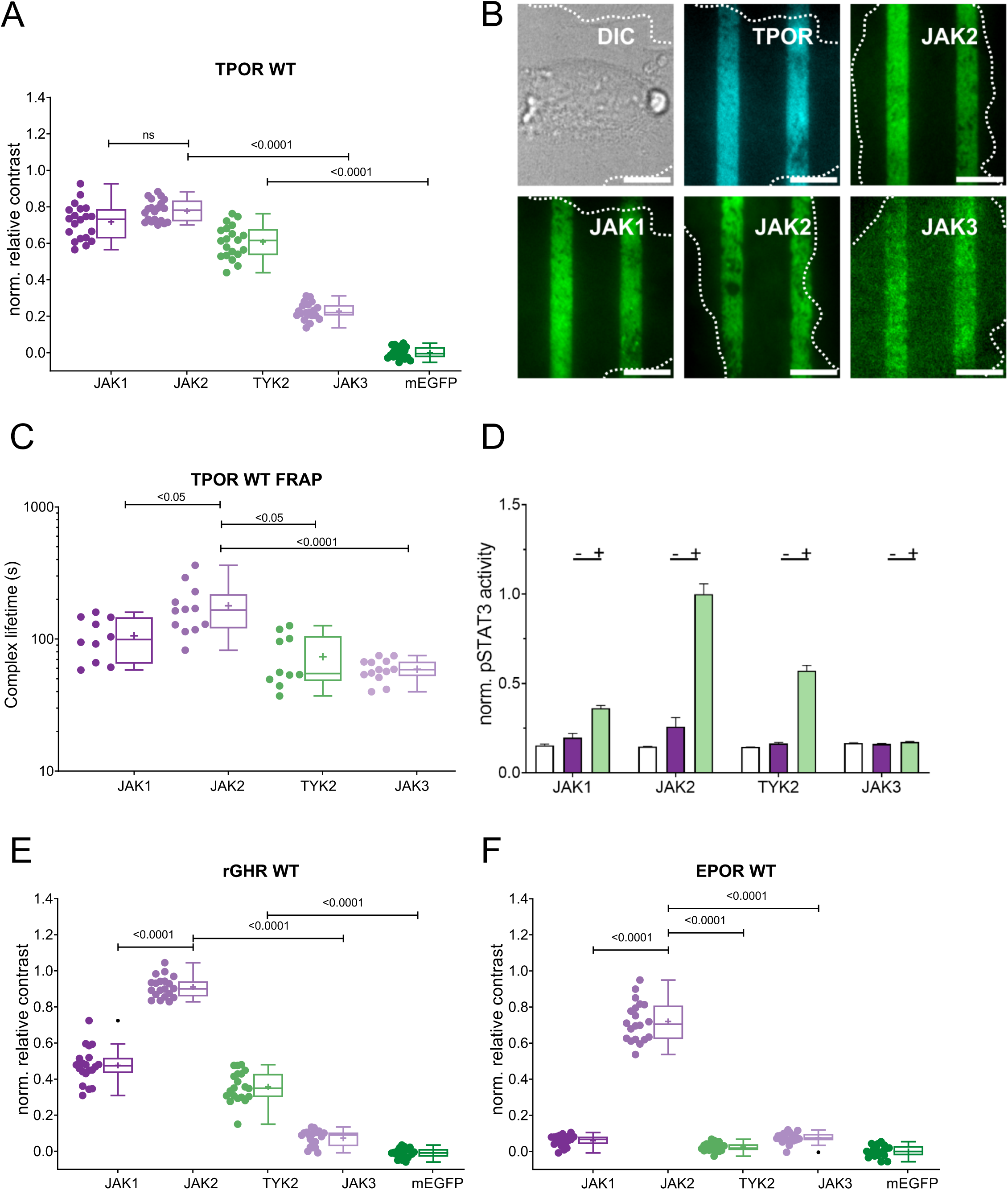
Promiscuous JAK binding to TpoR with differential signaling activity. **(A, B**) Binding of JAK1-FS, JAK2-FS, TYK2-FS and JAK3-FS to micropatterned HaloTag-mTagBFP-TPOR. Relative contrast of the JAK-FS-mEGFP channel (**A**) and representative images (**B**). (**C**) Comparison of the complex lifetimes obtained by fitting recovery curves with a single exponential. (**D**) Comparison of pSTAT3 levels in the absence (purple) and the presence of TPO (green) upon overexpression of JAKs FL. Untransfected cells are shown as negative control (white bars). (**E, F)** Binding of JAK1-FS, JAK2-FS, TYK2-FS and JAK3-FS to micropatterned HaloTag-mTagBFP-rGHR (**E**) or HaloTag-mTagBFP-EPOR (**F**). Relative contrast of the JAK-FS-mEGFP channel

For analyzing TPOR signaling activity in the presence of different JAKs, we turned to phospho-flow cytometry. To this end, HaloTag-mTagBFP-TPOR was co-expressed with different JAKs fused to mEGFP in WT HeLa cells and stimulated with TPO for 15 min. For phospho-flow analysis, cells were gated for high expression of the corresponding JAK, thus ensuring efficient competition with endogenous JAKs. Interestingly, we found highly different TPOR downstream signaling activity for different JAKs, despite the very similar recruitment to the receptor observed in live cell micropatterning experiments (Fig. 8D). Similar to TPOR and IFNARs, growth hormone receptor (GHR) shows promiscuity to the different JAKs (Fig. 8E). By contrast, erythropoietin receptor (EPOR) selectively recruited JAK2, while binding of other JAKs was negligible (Fig. 8F). These results highlight that beyond receptor binding, further constraints may control functional coupling of JAKs to the receptors.

## DISCUSSION

IFN-I signaling is activated by ligand binding the extracellular domains of its two-receptor chains, IFNAR1 and IFNAR2, resulting in the proximity of the ICD chains and associated JAK1 and TYK2 kinases, which cross-phosphorylate each other, the two receptor chains and transiently recruited STATs. The activated STATs drive the transcription of a large number of robust and tunable genes (*3*, *5*, *6*). Here, we re-visited this standard model using the AIR receptor system, which was shown to be an excellent tool for studying the structure/function of different cytokine receptors (*38*). The pattern of activation observed using the AIR system and IFN-β were similar, including STAT phosphorylation, gene expression, and antiviral activity. This suggests that the signal transmitted from the extracellular domain of IFNAR to the intracellular domain is not tightly dependent of the exact orientations of the domains or distances between the ECDs, rather than on the proximity of the two receptor chains, which drive JAK cross-phosphorylation (*62*).

We exploited the modularity of the AIR approach to test non-natural homodimeric pairing of IFNAR1 and IFNAR2, respectively. While AIR-dR1 did not promote activity, weak STAT1 and STAT2 phosphorylation was observed for AIR-dR2, resulting in activation of gene expression and partial antiviral protection. These results align with those reported previously (*41*). Wondering whether this effect is related to different pairing of JAK1 and TYK2, we probed by pull-down of AIR receptors which JAKs are associated with IFNAR1-ICD in the physiological context. In addition to TYK2, we found high levels of JAK1 being associated with IFNAR1. This surprising result led us to re-explore the roles of JAKs in IFNAR signaling. To obtain a non-biased view, we generated JAK1, JAK2 and TYK2 KO cells, and double JAK2/TYK2 KO cells. In line with previous studies, we confirmed that JAK1 is essential for IFN-1 signaling (*14*, *63*, *64*). However, we show that this requirement is not absolute, as TYK2 overexpression on the background of JAK1 KO produced a weak IFN-I signal. Likewise, partial IFN-I signaling activity observed on the background of TYK2 KO in HeLa cells was further elevated by overexpression of JAK1. The findings indicate that in addition to signaling through the typical JAK1/TYK2 heterodimer, the IFNAR complex is able to recruit JAK1/JAK1 and TYK2/TYK2 homodimers as well as JAK1/JAK2 heterodimers. However, these alternative dimer combinations exhibit diminished signaling potential compared to the canonical JAK1/TYK2 pairing. We confirmed promiscuous, competitive binding of different JAKs to both, IFNAR1 and IFNAR2 by live cell micropatterning. IFNAR1 was able to recruit TYK2, JAK1, and JAK2, despite JAK2 not being the canonical kinase for IFN-I signaling. Modeling JAK binding to IFNAR1 and 2 using AlphaFold further support the interchangeable binding of various JAK FERM domains to cytokine receptor ICDs. These results highlight that different JAKs compete for binding to IFNAR1 and IFNAR2 and that the receptorś actual choice of JAKs depend on their absolute and relative concentration in the cell. JAK1 is the dominantly expressed JAK member in HeLa cells (∼30-fold higher at protein level compared to TYK2) (*65*), which explains its efficient binding to IFNAR1 observed in pull-down experiments.

Despite binding IFNARs with similarly high efficiency, the signaling capacity of different JAK pairings is highly different. Reduced activation of IFN-I signaling can be explained by altered receptor trafficking that strongly depends on JAK identity and quantity (*66*, *67*), which was confirmed by our KO cell lines. Interestingly, we here found that in addition to TYK2, also JAK2 is involved in stabilizing IFNAR1 cell surface expression, either directly or indirectly. However, while JAK2 KO similarly reduced IFNAR1 cell surface expression levels as compared to TYK2, its impact on signaling activity was much milder. In line with this observation we found substantially higher capability to form ternary complexes in JAK2 KO as compared to TYK2 KO cells. Strikingly, IFNAR1 cell surface expression in TYK2 KO cells was compensated by JAK1 overexpression, but the recovery of signaling activity was minor. Taken together, these observations clearly implicate that different JAK homo- and heterodimers in the IFNAR complex can tune capacities of signaling activity from high (JAK1/TYK2) via medium (JAK1/JAK1 and JAK1/JAK2) to inactive (TYK2/TYK2) (Fig. 7B).

Systematic studies of receptor and JAK pairings based on a synthetic receptor approach has demonstrated that essentially all JAK combinations can be paired into a signaling-competent manner, yet the activity strongly depends of precise coupling between the transmembrane and the membrane-proximal JAK binding domain (*68*). In conjunction with our studies, these results suggest that the structural architecture of cytokine receptor dimers are optimized for certain combinations of JAKs. Given the highly different expression levels of different JAK members in different cell types (*68*), promiscuous JAK binding can be considered as a generic regulator of signaling activity. JAK binding studies with TPOR, GHR and EPOR as prototypic homodimeric cytokine receptors further support this hypothesis. High specificity was only observed for EPOR, while TPOR and GHR show similar promiscuity as IFNAR1 and IFNAR2. Yet, the potency of downstream signaling by TPOR showed striking differences for different JAK homodimers, which has previously also been related to differential orientational constraints required for functional pairing (*69*). Similarly, differential functional coupling of GP130 via different JAK homodimers has recently been described (*60*). We therefore expect that modulation of cytokine receptor activity by promiscuous JAK binding may apply relatively broadly, which can explain the pronounced pleiotropy common in this family (*5*, *16*, *48*, *60*, *66*, *67*, *70*). modulation of IFNAR signaling by differential JAK recruitments as characterized in this study opens a new perspective into the tunable nature of IFN-I signaling. IFNAR1 and IFNAR2 are ubiquitously expressed and the capability to respond against viral infection is vital for essentially all cell types of the body. Given the highly diverse expression of different JAKs, however, different JAK pairing and thus differential activity of IFN-I responses can be expected. Antiviral activity, however, is robustly maintained even in the absence of TYK2, which generally is expressed at much lower level as compared to JAK1. By contrast, tunable responses of IFN-I, such as antiproliferative and immunomodulatory activities require higher levels of TYK2, which indeed can be found in immune cells (*68*). Interestingly, type III interferon signaling, which is highly related to IFN-I signaling, but limited to certain cell types and antiviral activity, appears even less dependent on the presence of TYK2 (*47*).

In conclusion, this study provides valuable insights into the complex regulatory role of promiscuous JAK family members binding to cytokine receptors. Our findings open new perspectives for future research in cytokine receptor signaling and therapeutic interventions. JAK inhibitors are currently emerging as important drugs for diverse conditions, but the promiscuity of JAK recruitment is currently not being considered. Thus, the TYK2 inhibitor deucravacitinib will attenuate signaling of IFN-I and other cytokines in a much more cell-type specific manner than currently appreciated. Our findings substantiate that the combinatorial assembly of JAK kinases with cytokine receptor chains enables tailored signaling responses. This may allow the innate immune system to mount responses proportional to the severity of pathogens and stimuli encountered. Future work should explore whether dysregulated JAK-receptor promiscuity underlies aberrant cytokine signaling in autoimmunity and immunodeficiency. Elucidating the molecular grammar underlying differential JAK usage promises to uncover new therapeutic strategies for immunomodulation.

## MATERIALS AND METHODS

### Cell lines

HeLa cells, derived from a human cervical cancer cell line, were used for all experiments in this study. For culturing, Dulbecco’s Modified Eagle’s Medium (DMEM, Gibco 41965-039) supplemented with 10% fetal bovine serum (FBS, Gibco 12657-029), 1% pyruvate (Biological Industries 03-042-1B), and 1% penicillin-streptomycin (Biological Industries 03-031-1B) was used.

### AIR receptor construction and transfection

Transient transfections were carried out using JetPrime reagent (PolyPlus 114-07) following the manufacturer’s protocol. After 48 hours of transfection, cells were treated to assess AV activity, gene induction, and protein phosphorylation levels (WB).

### Generating KO cells using CRISPR-Cas9

All knockout (KO) cell lines in this study were generated using the CRISPR-Cas9 technology, as previously described (*14*, *17*). For the current study, we created a TYK2 knockout, JAK2 knockout, and the double TYK2/JAK2 knockout HeLa cell lines. To target the genes, we designed a single guide RNA (sgRNA) using the Benchling CRISPR Design Tool. The sgRNA sequence “5’-TGAATGACGTGGCATCACTG-3’” was designed to specifically target exon 6 of the TYK2 gene, while the sgRNA sequence “5’-CGTTGGTATTGCAGTGGCAG-3’” was designed to target exon 5 of the JAK2 gene. The sgRNA was cloned into the pX459 plasmid (Addgene plasmid #62988) and transfected into HeLa cells using JetPRIME (polyplus 114-07) following the manufacturer’s instructions. Clones were selected using puromycin resistance, and single-cells were expended and verified through Western blotting (WB) and genomic sequencing for the KO. Similarly, JAK1 KO and double IFNAR1/IFNAR2 knockout cell lines were generated using previously described methods (*14*, *17*).

### Protein expression and purification

The expression and purification of the AIR ligand was conducted following the procedures described (*71*). The plasmids pET28-14 His-bdSUMO-mEGFP-5aa-mCherry were introduced into T7 bacteria, which were then cultured in 2YT media supplemented with kanamycin. After the standard Ni-NTA purification process using Ni-NTA beads (Merck, cat. 70666-4) and subsequent Sumo protease digestion (concentration of sumo protease at 1:200), most of the proteins were loaded onto a Hi-trap Q HP anion exchange column (GE Healthcare, cat. 17115401). The final purification step involved using a size exclusion column, specifically Superdex 200 Increase 26/600 (GE Healthcare, cat. GE28-9893-34), to ensure thorough cleaning and separation of the proteins. IFN-α2 M148A (*72*) and IFN- α2 α8-tail (*73*) fused to an N-terminal ybbR-tag (*74*) were expressed in *E. coli* and refolded and purified from inclusion bodies as described previously (*75*). Site-specific labeling of the ybbR-tag via enzymatic phosphopantetheinyl transfer from CoA conjugated with DY 647P1 maleimide was carried out as described previously (*52*).

### WB analysis

For Western blot analysis, the cells were lysed in PBS (pH 7.4) containing 1% NP-40, 1 mM EDTA, and a cocktail of protease inhibitors (Sigma P8340), phosphatase inhibitor cocktail 2 (Sigma P5726), and phosphatase inhibitor cocktail 3 (Sigma P0044). The cell lysates were separated by 4-20% SDS-PAGE (GenScript M00657) and transferred to a nitrocellulose membrane (0.45 µm, Bio-Rad). For primary antibody incubation, the membranes were blocked with 5% bovine serum albumin (BSA). The blots were then probed with specific antibodies and detected using ECL substrate. The following antibodies were used: rabbit polyclonal anti-pSTAT1 (Tyr701) (Cell Signaling 9167S), rabbit polyclonal anti-PY-STAT2 (Tyr690) (Cell Signaling 4441S), rabbit monoclonal anti-pSTAT3 (Tyr705) (Cell Signaling Technology, 9145S), mouse monoclonal anti-pSTAT6 (pY641.18) (Santa Cruz Biotechnology Inc., SC-136019), mouse monoclonal anti-α-tubulin (Sigma-Aldrich, T9026), horseradish peroxidase (HRP)-conjugated anti-mouse antibody (Jackson ImmunoResearch, 115035146), and HRP-conjugated anti-rabbit antibody (Jackson ImmunoResearch, 111035144). The Western blots were imaged using an Odyssey Fc imager (LI-COR Biosciences), and quantitative analysis was performed using Image Studio Lite software (LI-COR Biosciences).

### Flow cytometry

Surface levels of IFNARs were determined by flow cytometry according to the methodology described in references (*5*, *14*, *17*). Mouse monoclonal antibodies, AA3 and 117.7 (*76*), were employed, specifically recognizing IFNAR1 and IFNAR2, respectively. Adherent cells were initially detached using phosphate-buffered saline (PBS) supplemented with 5 mM EDTA. Subsequently, the cells were resuspended in PBS containing 3% fetal bovine serum (FBS, Gibco 12657-029) and 0.5 mM EDTA. The first antibody was added, followed by signal amplification using biotinylated rat anti-mouse IgG (Jackson Immunochemicals) and streptavidin-allophycocyanin (APC) (BD Biosciences). Propidium Iodide (Sigma-Aldrich p-4864) was used to gate live cells. Samples were analyzed using the CytoFLEX S flow cytometer, and the median of the APC signal was extracted.

### Phospho-flow cytometry

Cells were cultured in DMEM supplemented with 10% FBS, 1% pyruvate, and 1% penicillin-streptomycin. Cells were seeded in 15cm plates at a density of 1-2 million cells per plate. The next day, cells were washed once with 10mL of pre-warmed 5 mM EDTA in DMEM and incubated with 10 mL of EDTA at 37°C for 5-10 minutes until cells detached. Cells were collected, centrifuged at 500g for 3 minutes, and resuspended in EDTA at a density of 1 million cells per 80 µL. 80 µL of cell suspension was aliquoted into each well of a 96-well plate. Cells were treated with cytokines diluted in EDTA and incubated at 37°C for 15-240 minutes.

After cytokine stimulation, cells were fixed by adding 100µL of 4% paraformaldehyde (PFA) directly to the wells, mixing thoroughly, and incubating at room temperature for 10 minutes. Plates were centrifuged at 500g for 2 minutes and the supernatant carefully removed. Cells were quenched with 200 µL of 2M glycine for 15 minutes. After quenching, cells were washed twice with 200 µL cold methanol. Cells were then permeabilized by resuspension in 130µL cold methanol and incubated for 20 minutes at room temperature. Next, 130 µL of barcoding dye (Pacific Blue and AlexaFluor-488) was added to each well, mixed, and incubated at room temperature in the dark for 20 minutes. Cells were washed twice with methanol to remove excess dye. Finally, cells were washed with 1% BSA in PBS. After barcoding, cells were aliquoted into a 96-well plate at 80 µL per well. Fluorophore-conjugated antibodies were added at optimal dilutions and incubated for 30 minutes at room temperature in the dark. Cells were washed twice with 1% BSA/PBS, resuspended in 200 µL, filtered, and analyzed by flow cytometry. Compensation was performed using single stained samples to account for spillover of Pacific Blue and AlexaFluor-488 into the APC channel. (Antibody: pSTAT1: BD Phosflow 612597, pSTAT2: cell signaling 907405, pSTAT3: BD Phosflow 557815, pSTAT5: BD Phosflow 612599, pSTAT6: BD Phosflow 612601).

pSTAT activation by TPOR in the presence of different JAKs was probed by phospho-flow cytometry as described previously(*53*). To this end, JAK1 KO HeLa cells were co-transfected with the TPOR fused to HaloTag-mTagBFP and corresponding JAK fused to mEGFP. Cells were simultaneously stimulated for 15 min at 37°C with the corresponding ligand followed by PFA fixation (4%) for 15 min at RT. After fixation, cells were centrifuged at 300g for 5 min at 4°C. Cell pellets were resuspended and permeabilized in ice-cold methanol and kept for 30 min on ice. After permeabilization, tyrosine-phosphorylated STAT3 (pY705) was stained using a murine monoclonal antibody labeled with AlexaFluor 647 (Cell Signaling #4324, 1:100 dilution). Cells were analyzed on a flow cytometer (Cytoflex S, Beckman Coulter) and gated for receptor and JAK expression levels based on the intensity in the BFP and GFP channel, respectively. Median fluorescence intensity (MFI) of pSTAT3 was measured for stimulated and non-stimulated cells. Non-transfected cells were used as a control.

### Quantitative PCR analysis

As described previously (*5*, *14*, *17*), the expression levels of selected human interferon-stimulated genes were assessed using the Applied Biosystems ViiA 7 Real-Time PCR System. Fast SYBR Green Master Mix (Applied Biosystems) was utilized for the PCR reactions, and cDNA samples were generated from 1 µg of total RNA using the high-capacity cDNA reverse transcription kit (Applied Biosystems). The RNA was extracted using the NucleoSpin RNA kit (Macherey-Nagel). For quantitative PCR (qPCR), 14.5 ng of cDNA was used in a total reaction volume of 7 µL with primers as listed in Table S2. To determine the relative expression levels, the ΔΔCT (cycle threshold) method was employed, where the fold change was calculated using the formula RQ = 2^(-ΔΔCT). The endogenous control gene Hypoxanthine-guanine phosphoribosyltransferase1 (HPRT1) was used as the reference gene, with its expression levels measured in the same sample.

### MARseq

The cells were treated with either 2 nM IFN-β or AIR ligand for 16 hrs or 6 hrs. The RNA isolation procedure employed to purify the RNA is as for qPCR. The MARS-seq experiment was conducted at the INCPM units at the Weizmann Institute of Science. This experimental protocol involved barcoding the samples through reverse transcription using an oligo dT primer, followed by pooling of the samples. Subsequent molecular reactions were performed to amplify and prepare specimens for Illumina sequencing(*77*, *78*). To analyze the MARS-seq data, the UTAP transcriptome analysis pipeline was utilized(*79*).

### Antiviral (AV) and antiproliferative (AP) assays

1.2 × 10^4^ or 2 × 10^3^ cells were grown overnight on flat-bottomed 96-well plates for the AV and AP assays respectively. Cells were treated with ten 3-fold IFN-β or AIR ligand dilutions starting from 500 pM or 50 nM for AV and AP, respectively. To assess AV protection against VSV and EMCV, the extent of cytopathic effect inhibition caused by the viruses were determined. Four hours after the addition of treatment, VSV or EMCV were introduced and incubated for 18 or 20 hours, respectively. AP activity was measured 96 hours after the addition of IFN-β. Cell viability was determined using crystal violet staining, as previously described. The EC50 values and cell sensitivity to the indicated ligands were determined by analyzing the response curve implemented in the GraphPad Prism software package version 9.5.0.730.

### Cell micropatterning and image analysis

The binding of JAKs to the receptors in the plasma membrane was quantified by live cell micropatterning using the HaloTag as previously described (*80–82*). Surfaces functionalized with the micropatterned HaloTag ligand (HTL) were fabricated by microcontact printing, as described in detail previously (*83*). Standard glass coverslips for fluorescence microscopy were cleaned in a plasma cleaner for 10 min. PDMS stamps were inked with 0.5 mg/mL poly-L-lysine-graft-poly (ethylene glycol) (PLL-PEG) conjugated with the HTL (PLL-PEG-HTL) in PBS buffer for 10 min and then placed with additional weight onto the glass coverslips for 10 min to generate HTL micropatterns. After removing the stamps, the coverslips were incubated with a 4:1 mixture of 0.5 mg/mL of methoxy-terminated PLL-PEG (PLL-PEG-MeO) and 0.5 mg/mL PLL-PEG conjugated with the peptide RGD (PLL-PEG-RGD)(*81*) in PBS buffer for 2 min to backfill the uncoated area to allow cell adhesion. The surface was then rinsed in MilliQ water and dried with nitrogen. Cells were transiently transfected with receptor N-terminally fused to the HaloTag and mTagBFP (HaloTag-mTagBFP-IFNARs), truncated JAK1/JAK2 and TYK2 comprising only the FERM and SH2-like domains C-terminally linked to mEGFP (JAK-FS). Transfected cells were transferred onto micropatterned surfaces 24-36 h post transfection and cultured for 15-20 h with a penicillin and streptomycin (PAA) medium. Micropatterned cells were then imaged by TIRF microscopy using an inverted microscope (Olympus IX81) equipped with a 4-line TIRF condenser (Olympus TIRF 4-Line LCl), a CMOS camera (ORCA-Flash 4.0, Hamamatsu) and lasers at 405 nm (100 mW), 488 nm (150 mW). A 60× and a 100× objective with a numerical aperture of 1.49 (UAPON 60×/1.49, Olympus, UAPON 100×/1.49) were used for TIRF excitation. The excitation beam was reflected into the objective by a quad-band dichroic mirror (zt405/488/561/640rpc), and the fluorescence was detected through a quadband pass filter (BrightLine HC 446/523/500/677). For multicolor experiments, a fast emission filter wheel equipped with suitable emission filters (BrightLine HC 445/45, BrightLine HC 525/50, BrightLine HC 600/37, and BrightLine HC 697/58) was utilized to avoid spectral cross-talk. Data acquisition was performed with the acquisition software Olympus CellSens 2.2.

Image analysis and image processing were performed using ImageJ/ Fiji (NIH, Bethesda, MD). Image processing comprised cropping, scaling, rotation, and adjusting brightness and contrast levels. The fluorescence contrast of the bait proteins (*C_bait_*) was calculated as follows:

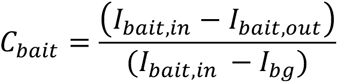

*I_bait, in,_* and *I_bait, out_* denote the mean pixel intensities in the mtagBFP channel from selected ROIs inside and outside, respectively, of the HTL-functionalized areas. *I_bg_* is the background intensity from the glass surface obtained from an ROI outside the region covered by the cells. The contrast of the prey proteins *C_prey_* was accordingly obtained from the background corrected mean pixel intensities from selected ROIs inside and outside HTL-functionalized areas:

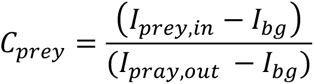

with *I_prey, in,_* and *I_prey, out_* denoting the mean pixel intensities in GFP channel from selected ROIs inside and outside, respectively, of the HTL-functionalized areas. Since *C_prey_* is dependent on *C_bait_*, *C_prey_* was corrected for variances in *C_bait_* by dividing *C_prey_* by *C_bait_*:

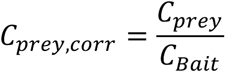

The contrast values were normalized by subtracting from all matters of one set of experiments the mean of the negative control and dividing all values of one set of investigations by the positive control standard. For statistical analysis, 20 to 30 cells were analyzed, and the calculated contrast values were visualized in a box plot. Two-sample Kolmogorov-Smirnov-Tests were performed to calculate statistical significance.

### Single-molecule binding assays

Single-molecule imaging was carried out by total internal reflection fluorescence microscopy (TIRFM) using an inverted microscope (IX83-P2ZF, Olympus) equipped with a motorized quad-line TIR illumination condenser (cellTIRF-4-Line, Olympus) as recently described in detail(*84*). DY647 was excited using a 642 nm laser (2RU-VFL-P-500-642-B1R, MPB Communications). Fluorescence was filtered by a penta-band polychroic mirror (zt405/488/561/640/730rpc, Semrock) and excitation light was blocked by a penta-band bandpass emission filter (BrightLine HC 440/521/607/694/809, Semrock). HeLa cells were cultivated at 37°C under 5% CO_2_ in MEM with Earle’s salts supplemented with 10% FBS superior (Merck KGaA), 2 mM L-Alanyl-L-Glutamine (Biochrom), 1% non-essential amino acids (Merck KGaA) and 10 mM HEPES buffer (Carl Roth). Cells were transfected with single plasmids at 60-70% confluency by PEI for 4 hours, followed by medium exchange and regeneration for 16-24 hours. The day before microscopy, cells were detached by room-temperature treatment of Accutase (Innovative Cell Technologies) and seeded on microscopy cover slides coated with a 50/50 (w/w) mixture of poly-L-lysine graft copolymers of polyethylene glycol (PLL-PEG) that were modified with an RGD-peptide and a terminal methoxy group, respectively (*83*, *85*). Imaging experiments were performed at 25°C in phenol red-free DMEM medium supplemented with 10% FBS in presence of an oxygen-scavenging system composed of glucose oxidase (4.5 U*mL^-1^), catalase (540 U*mL^-1^), glucose (4.5 mg*mL^-1^), ascorbic acid (1 mM) and methyl viologen (1 mM) (*86*) to increase photostability.

5 nM of either IFN-α2 M148A or IFN-α2 α8-tail were added to the cells and kept in solution throughout the experiment. After 5 min of incubation videos of viable cells were recorded at 30 fps for typically 150 consecutive frames using CellSens 2.2 (Olympus) as acquisition software. Microscopy image stacks were subjected to single-molecule localization analysis using a custom-made MATLAB script called SLIMfast 4C as described with more detail in (*84*). Surface densities were calculated as mean number of detected mobile molecules/frame per cell. Only single molecule events that were tracked for a minimum of 10 consecutive frames (320ms) and travelled at least 100 nm in their course were considered mobile molecules. Statistical analysis was performed from typically 15 cells recorded for each condition. Two-sample Kolmogorov-Smirnov-Tests were performed to calculate statistical significance.

### FRAP measurements

The interaction stability was determined by fluorescence recovery after photobleaching (FRAP). A rectangular region of interest (ROI) within the bleached area of the pattern and a rectangular ROI within the bleached area but outside the patterned area was chosen for obtaining intensity values per pixel over time, respectively. Corrected FRAP curves were determined using the following equation:

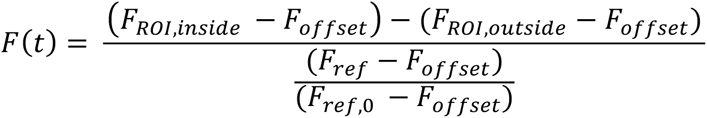

With *F_ROI, inside_* and *F_ROI, outside_* being the fluorescence intensities inside and outside the pattern, respectively, within the bleached spot. *F_ref_* is the fluorescence intensity of an unbleached ROI inside the micropattern, and *F_ref_*, *_0_* is the fluorescence intensity of this ROI before the bleaching experiment. *F_ref_* was used as a normalization factor to correct photobleaching during FRAP experiments. The offset intensity (*F_offset_*) was determined from an ROI outside the cell and subtracted from all intensity values. Image analysis to obtain corrected FRAP curves was performed using a MATLAB script. To obtain lifetime values, the corrected FRAP curves F(t) were fitted by a monoexponential function.

## Supplementary Materials

Video S1. Single molecule TIRF microscopy of fluorescent IFNα2 M148A/α8tail binding to endogenous IFNAR subunits on the surface of WT HeLa cells.

## Acknowledgments

We sincerely thank Dr. Moshe Goldsmith for his invaluable assistance and dedicated maintenance of the flow cytometry instrument in the Department of Biomolecular Science at the Weizmann Institute of Science. We are also grateful to the INCPM for conducting the MARSseq and the MS experiments described in this manuscript, Specially to Dr. Amir Prior. Furthermore, we sincerely thank Prof. Jürgen Scheller’s lab for generously sharing the plasmids containing the sequences for the high-affinity GFP and mCherry nanobodies. We thank G. Hikade and H. Kenneweg technical assistance.

## Funding

This work was supported by grants from:

Minerva Foundation number 714144 (GS)

Deutsche Forschungsgemeinschaft SFB 1557/P13 (JP)

Lower Saxony – Israel grant number A128369 (JP and GS)

## Author contributions

Conceptualization: EZ, TM, JP, GS

Methodology: EZ, TM, JSB, BS, JP, GS

Investigation: EZ, TM, JSB, BS, JP, GS

Visualization: EZ, TM, JSB, BS, JP, GS

Funding acquisition: JP, GS

Project administration: JP, GS

Supervision: JP, GS

Writing – original draft: EZ, JP, GS

Writing – review & editing: EZ, TM, JSB JP, GS

## Data and materials availability

All data needed to evaluate the conclusions in the paper are present in the paper or the Supplementary Materials. All materials can be shared through a material transfer agreement.

## Supplementary materials

**Table S1.** Mass spectrometry analysis of the pull-down AIR-R1.

| Description | Coverage (%) | Unique Peptides | Peptides | PSMs |
| --- | --- | --- | --- | --- |
| JAK1 HUMAN | 67 | 89 | 91 | 1859 |
| TYK2 HUMAN | 54 | 55 | 57 | 669 |
| K2C1 HUMAN | 54 | 38 | 42 | 365 |
| K1C10 HUMAN | 47 | 28 | 34 | 328 |
| AIR Ligand-GFP-mCherry | 69 | 27 | 27 | 305 |
| K22E HUMAN | 75 | 37 | 43 | 233 |
| TRYP PIG | 25 | 4 | 4 | 230 |
| K1C9 HUMAN | 60 | 28 | 29 | 204 |
| DHE3 BOVIN | 29 | 15 | 15 | 82 |
| AIR- R1 | 40 | 7 | 7 | 64 |
| PRDX1 HUMAN | 20 | 5 | 5 | 39 |
| K1C15 SHEEP | 13 | 1 | 6 | 39 |
| SYH HUMAN | 8 | 4 | 4 | 21 |
| UBIQ HUMAN | 41 | 3 | 3 | 13 |
| CATA HUMAN | 11 | 4 | 4 | 12 |
| KRHB4 HUMAN | 8 | 2 | 5 | 12 |
| K1H2 HUMAN | 5 | 1 | 2 | 10 |
| TRFE HUMAN | 4 | 2 | 2 | 5 |
| K2M2 SHEEP | 8 | 2 | 3 | 5 |
| THIO HUMAN | 21 | 2 | 2 | 4 |
| KRHB5 HUMAN | 5 | 1 | 2 | 4 |
| NEDD8 HUMAN | 17 | 1 | 1 | 2 |

**Table S2.**
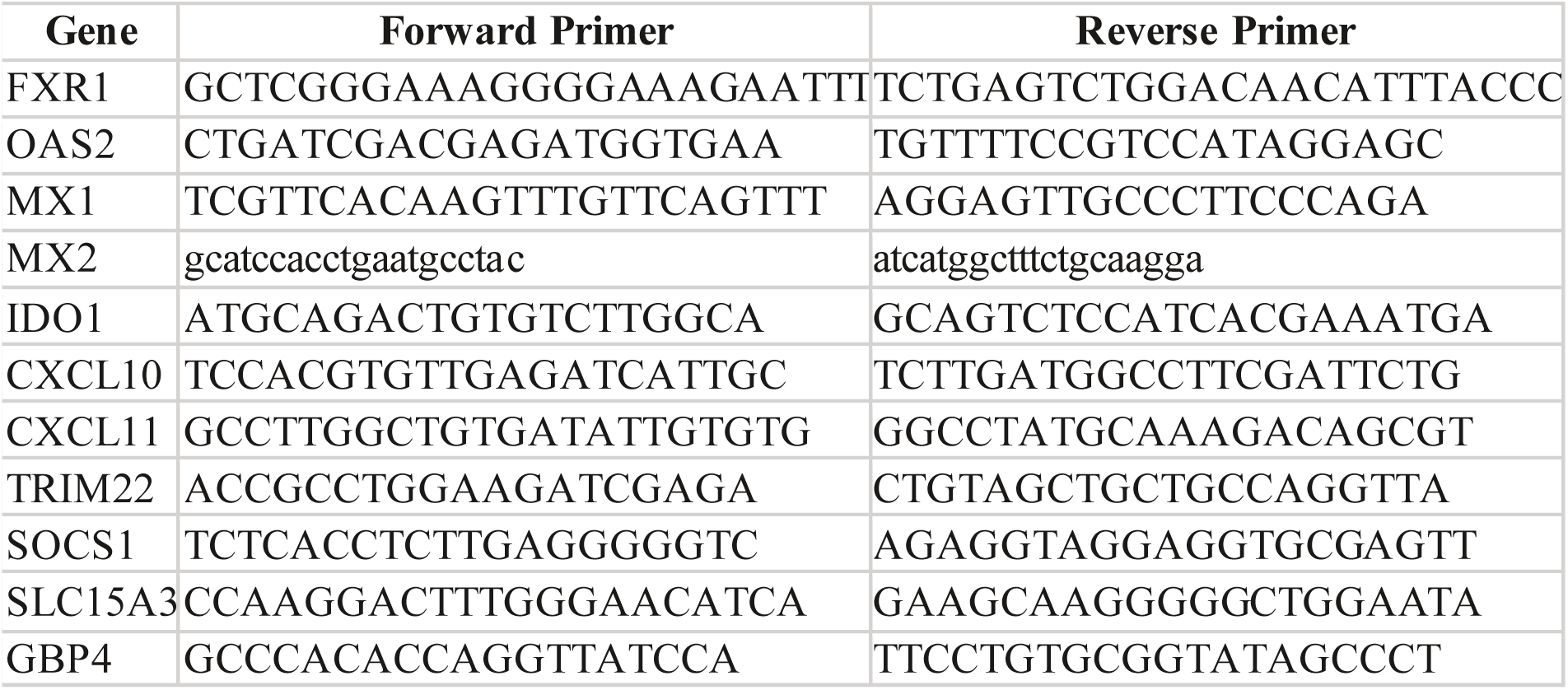
Primers for real-time PCR.

**Figure S1.**
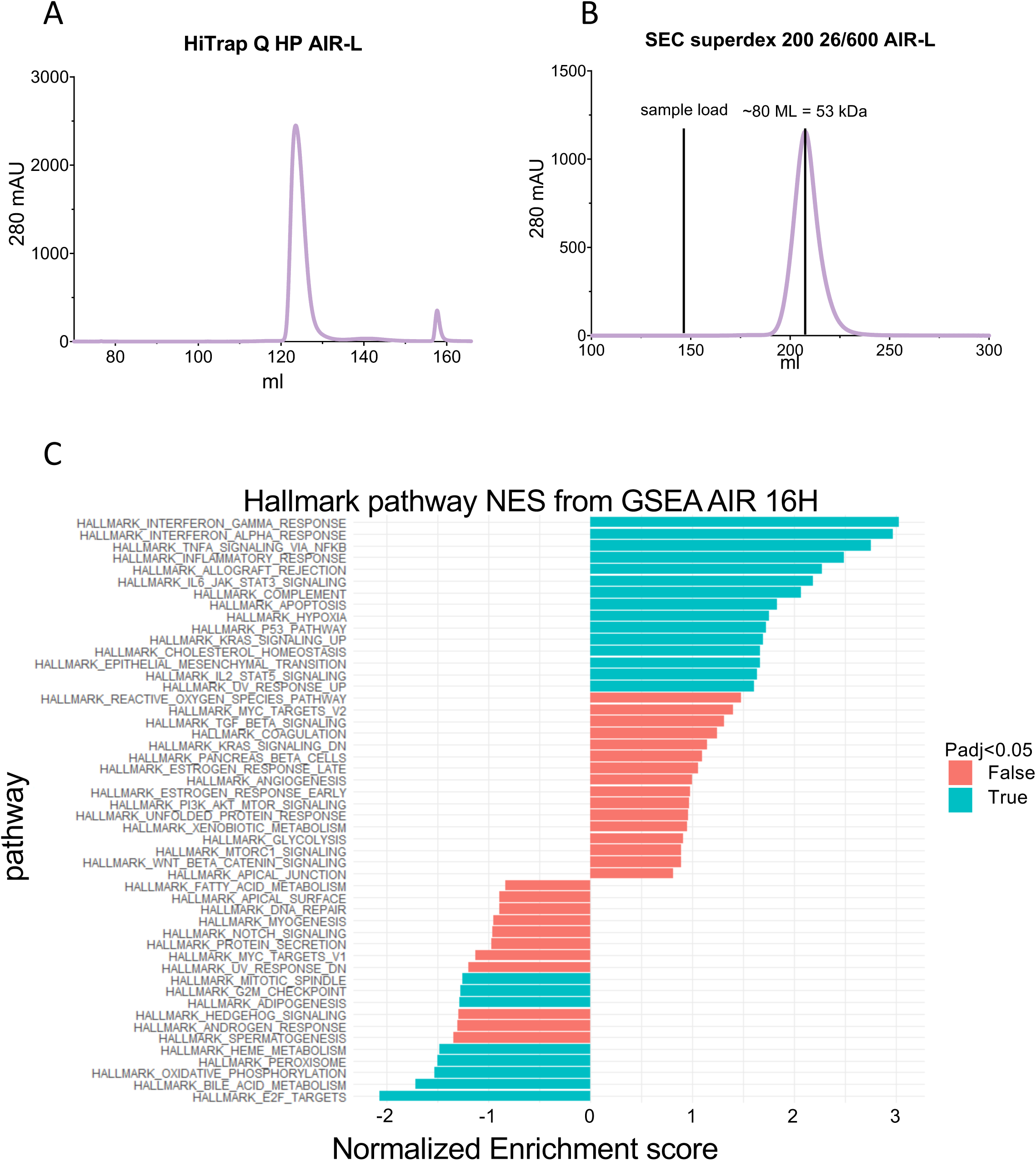
AIR ligand purification and GSEA analysis. AIR ligand was purified using using Ni-NTA beads and sumo cleavage, followed HiTrap Q HP anion exchange chromatography, with the protein being detected at 280 nm (mAU) (**A**). For final purification, size exclusion chromatography (superdex 200 26/600 chromatography detected at 280 nm (mAU) was used (**B**). (**C**) Gene Set Enrichment Analysis (GSEA) of MARS-Seq AIR treated for 16H with AIR-L. Pathways marked in pale blue have a p-adjusted value under 0.05.

**Figure S2.**
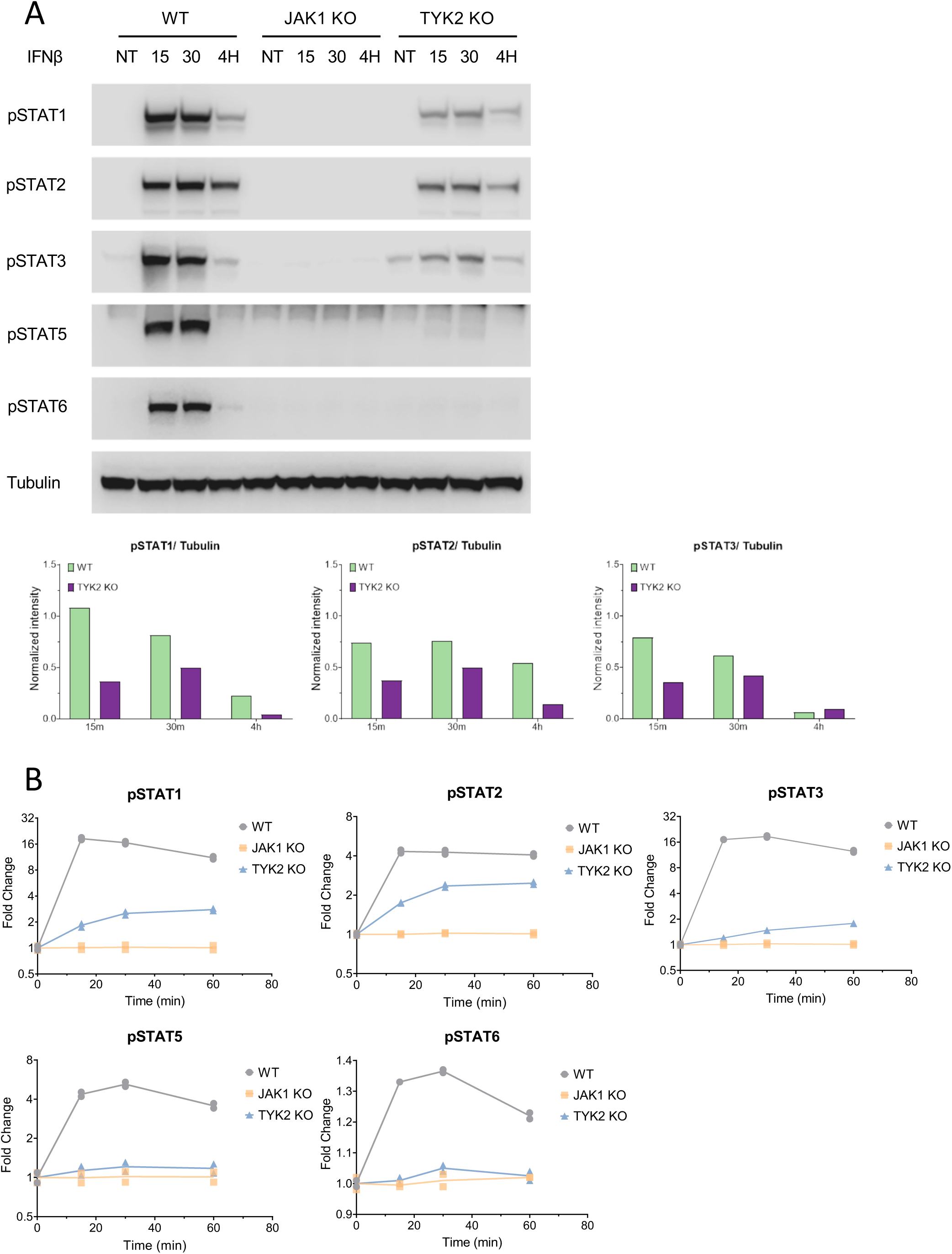
Phosphorylation of STATs in TYK2 and JAK1 KO cells. (**A**) HeLa (WT), JAK1 KO and, TYK2 KO cells were treated with 2 nM IFN-β for15, 30 min and 4 hours, and analyzed by Western blotting for phosphorylated STAT proteins. Tubulin is a loading control. **B**) Phospho-flow analysis of STAT phosphorylation at different times after induction with 1 nM. The data is one example out of three biological repeats.

**Figure S3.**
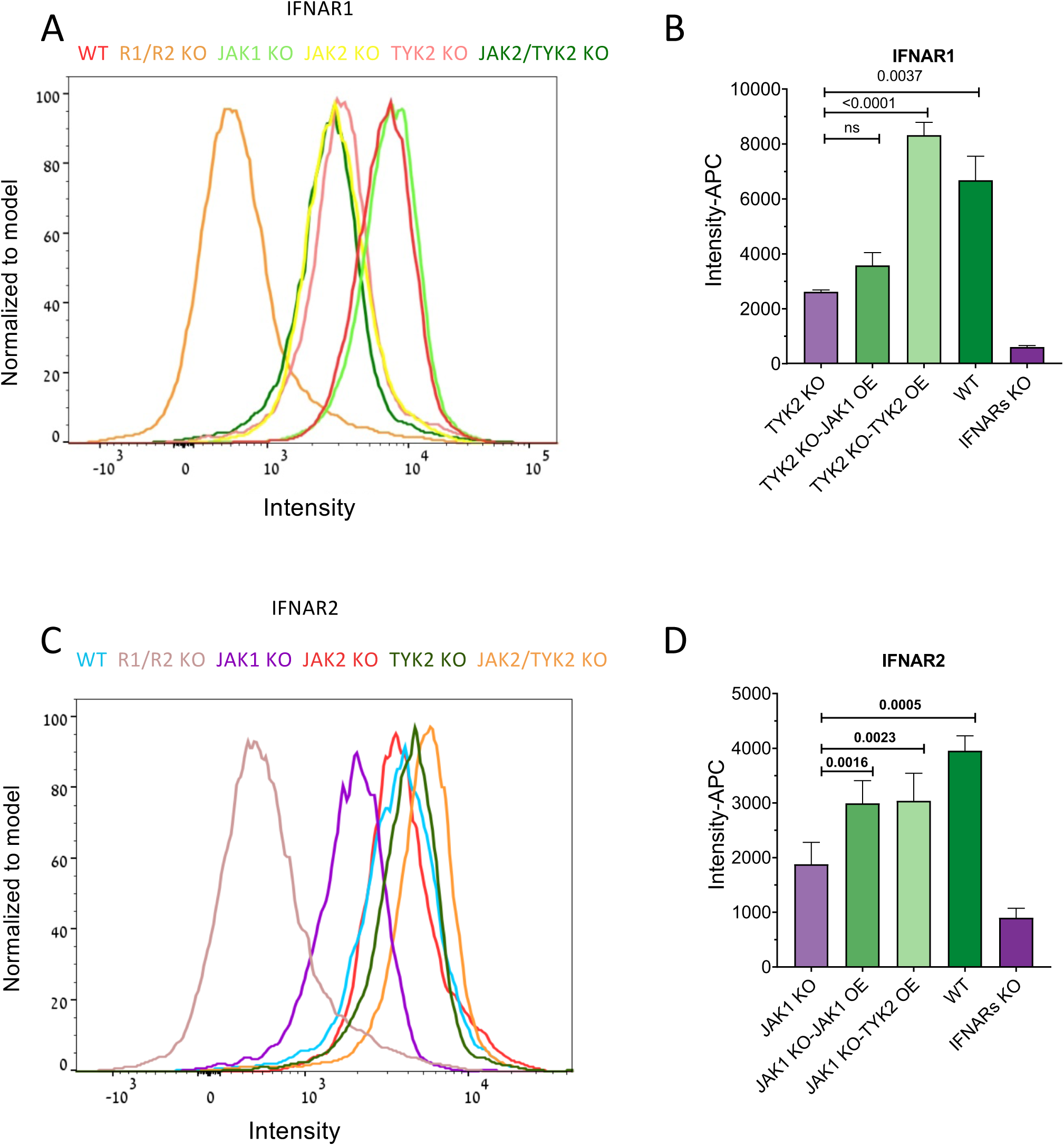
Cell surface receptor expression levels is regulated by JAKs. (A-D) Quantification of receptor cell surface expression levels in HeLa WT and KO cells by flow cytometry using antibodies against IFNAR1 (A and, B) and IFNAR2 (C and, D). OE stands for over expression.

**Figure S4.**
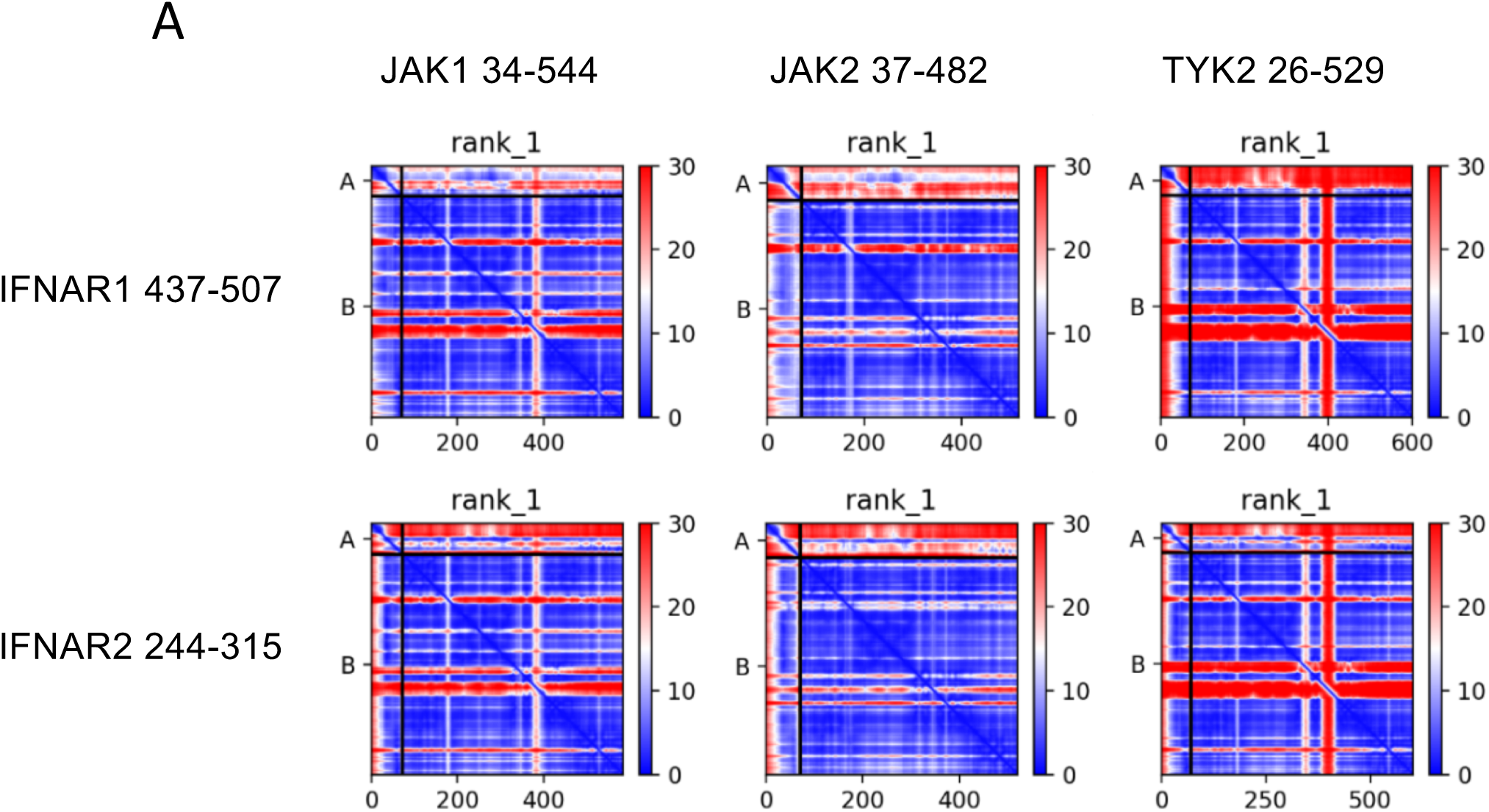
AlphaFold score. Predicted aligned error (page) for the six different model of the JAKs (FERM and SH2 domain) and the IFNARs ICD receptor presented in Fig. 7A.

